# Synthetic antigen-presenting cells reveal the diversity and functional specialisation of extracellular vesicles composing the fourth signal of T cell immunological synapses

**DOI:** 10.1101/2021.05.29.445691

**Authors:** Pablo F. Céspedes, Ashwin Jainarayanan, Lola Fernández-Messina, David G. Saliba, Salvatore Valvo, Audun Kvalvaag, Lina Chen, Elke Kurz, Charity Ganskow, Huw Colin-York, Marco Fritzsche, Yanchun Peng, Tao Dong, Errin Johnson, Jesús A. Siller-Farfán, Omer Dushek, Erdinc Sezgin, Ben Peacock, Alice Law, Dimitri Aubert, Simon Engledow, Moustafa Attar, Svenja Hester, Roman Fischer, Francisco Sánchez-Madrid, Michael L. Dustin

## Abstract

The T cell Immunological Synapse (IS) is a pivotal hub for the regulation of adaptive immunity by endowing the exchange of information between cells engaged in physical contacts. Beyond the integration of antigen (signal one), co-stimulation (signal two), and cytokines (signal three), the IS facilitates the delivery of T-cell effector assemblies including supramolecular attack particles (SMAPs) and extracellular vesicles (EVs). How these particulate outputs differ among T -cell subsets and how subcellular compartments and signals exchanged at the synapse contribute to their composition is not fully understood. Here we harnessed bead-supported lipid bilayers (BSLBs) as a tailorable and versatile technology for the study of synaptic particle biogenesis and composition in different T-cell subsets, including CART. These synthetic antigen-presenting cells (APCs) facilitated the characterisation of trans-synaptic vesicles (tSV) as a heterogeneous population of EVs comprising among others PM-derived synaptic ectosomes and CD63^+^ exosomes. We harnessed BSLB to unveil the factors influencing the vesicular release of CD40L, as a model effector, identifying CD40 trans presentation, T-cell activation, ESCRT upregulation/recruitment, antigen density/potency, co-repression by PD-1 ligands, and its processing by ADAM10 as major determinants. Further, BSLB made possible the comparison of microRNA (miR) species associated with tSV and steadily released EVs. Altogether, our data provide evidence for a higher specialisation of tSV which are enriched not only in effector immune receptors but also in miR and RNA-binding proteins. Considering the molecular uniqueness and functional complexity of the tSV output, which is also accompanied by SMAPs, we propose their classification as signal four.

**Graphical abstract:** 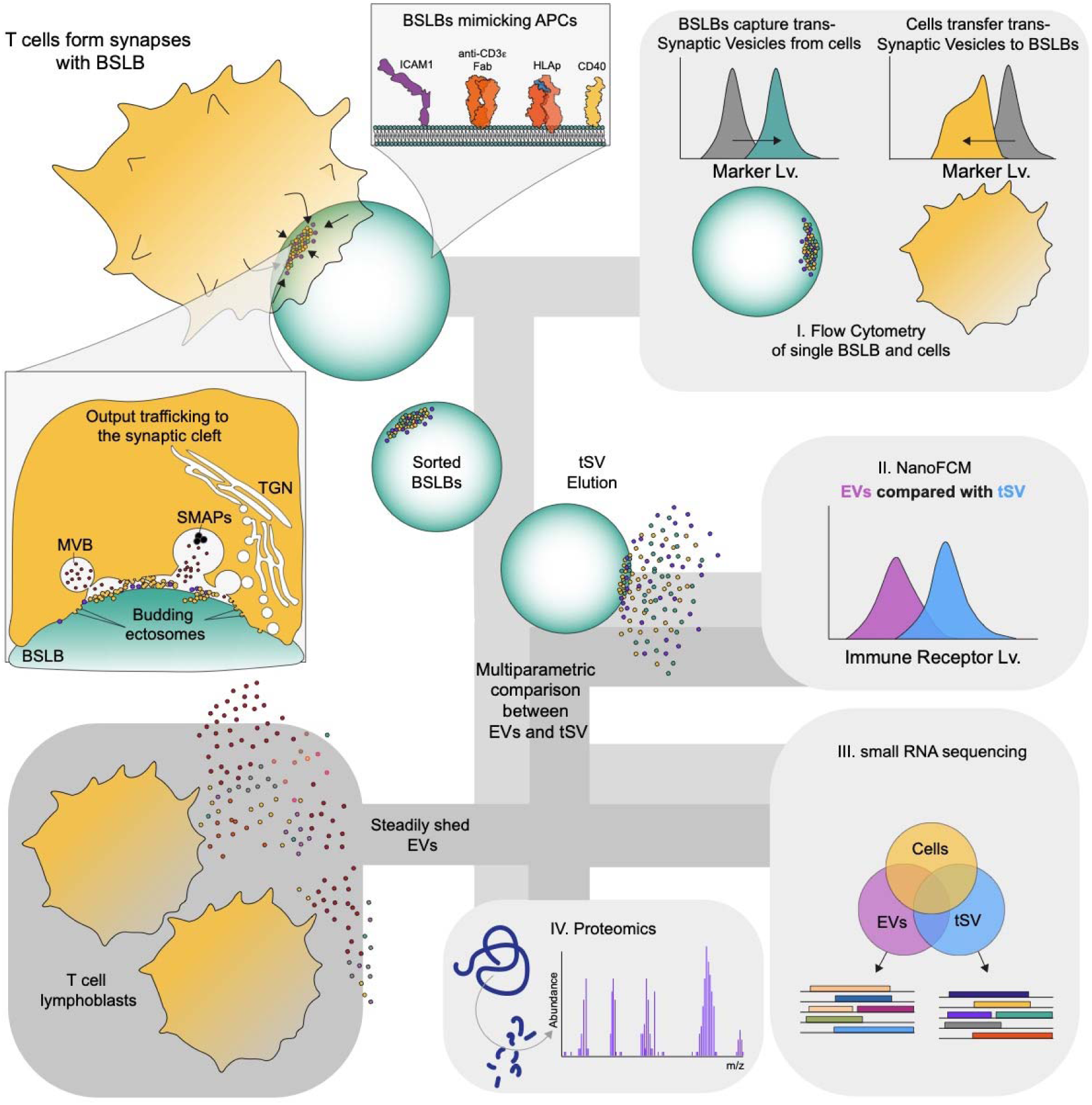

**Highlights:**

- Bead Supported Lipid Bilayers (BSLB) reconstituting antigen-presenting cells support synapse assembly by T cells and the release of effector particles.
- BSLB facilitate the dissection of the cellular machineries and synapse composition shaping the released tSV.
- tSV and their steadily released counterparts have a different composition. TSV show a higher enrichment of effectors including immune receptors, miR, RNA- and other nucleic acid-binding proteins, than EVs.

## INTRODUCTION

T lymphocytes are key players in the regulation and promotion of adaptive immunity. Helper T cells (TH), cytotoxic T lymphocytes (CTL) and regulatory T cells (Treg) shape cellular networks through the assembly of cell-cell interfaces termed Immunological Synapses (IS) with antigen-presenting cells (APCs) and other T cells[1, 2]. These ISs enable three key receptor-ligand driven signals critical to mounting an immune response or maintaining self-tolerance: 1) antigen recognition, 2) co-stimulation and co-repression, and 3) sensing of soluble cytokines released into the synaptic cleft. Here we propose a fourth type of trans-synaptic signal based on trans-synaptic supramolecular effectors and provide a general tool to study these challenging to isolate signaling entities.

In the past 10 years, the IS has been recognized as a site for bidirectional relay of supramolecular effectors, including polarized exosomes (PE) and supramolecular attack particles (SMAPs) released from multivesicular bodies[3] into the synaptic cleft, synaptic ectosome (SE) budding from the T cell plasma membrane (PM) across the synaptic cleft and trans-endocytosis of PM fragments across the synaptic cleft [4–6]. PE and SE are actively formed by the donor T cells through action of the endosomal sorting complexes for transport (ESCRT) machinery and together make up the trans-synaptic vesicles (tSV). Trans-endocytosis and trogocytosis result in taking of small or large, respectively, membrane fragments from one cell into another [7–11]. SMAPs are similar in size to exosomes and SE but lack a phospholipid membrane and instead have a core-shell structure that enables transfer of complex cargo[6]. tSVs and SMAPs have been studies by capture on synthetic APC, but the ability to manipulate them after capture has been limited and we offer a new tool through this resource.

While tSV, trans-endocytosed fragments and SMAPs are consumed by the synaptic partner, extracellular vesicles (EVs) formed by the ESCRT machinery and steadily released into the media can be collected from the culture supernatant of activated T cells[12]. Previous work has used EVs as a surrogate for tSV. While this approach has had some predictive power, a comparison of tSV and EVs is critical for progress in the field of cell-cell communication. Here, we develop a resource to collect tSV released from three major types of T cells and compared tSV and sEV released from the same cells. The approach should be applicable to study tSV generated in other settings.

## RESULTS

### Trans-synaptic vesicles are larger and carry more cargo than sEV

Supported lipid bilayer (SLB) on glass substrates have been used extensively to study cell recognition processes between live cells and SLB presenting molecular components from a natural interaction partner. SLB on planar supports presenting surrogate antigen, adhesion and co-stimulatory receptors were first used to characterize tSV and SMAPs from helper T cells and cytotoxic T cells, respectively. The advantage of the SLB is that following T cell release of tSV or SMAPs, the respective T cells can be selectively released, leaving the intact tSV or SMAPs behind for analysis by imaging or mass spectrometry. SLB can be readily formed on 5.0 ± 0.05 µm glass beads (BSLB) to improve compositional analysis [5]. We have further improved on this first generation of BSLB to develop a general resource for study of tSV and SMAPs in any synaptic model.

Our second generation BSLB model uses a surrogate antigen, a 14His tagged α-CD3ε Fab, that can be gently eluted from the SLB to enable release of tSV. We used fluorescence correlation spectroscopy to demonstrate that the recombinant α-CD3ε Fab and a co-incorporated Abberior Star-Red phosphoethanolamine had diffusion coefficients of 0.68 ± 0.39 and 3.2 ± 0.79 µm^2^/s, respectively, which are comparable to ranges reported for PM proteins and lipids [13–15]. To analyze tSV from helper T cells we used BSLB presenting a range of α-CD3ε Fab densities, and physiological densities of the adhesion molecule ICAM1 (200 molec./µm^2^) and costimulator receptor CD40 (20 molec./µm^2^), which are characteristic of natural APCs. Time-lapse imaging of T cell-BSLB co-cultures demonstrated that BSLB instigated transfer of CD40L from stimulated T cells, which left a “synaptic stamp” on the engaged BSLBs composed of the transferred tSV (Fig. 1A). To enable the synchronous release of BSLBs from conjugates, we gradually cooled the cell-BSLB co-culture to 4°C to promote dissociation of the LFA1-ICAM1 interaction while minimizing mechanical stress on BSLB and their tethered tSV. Following this treatment, flow cytometry (FCM) revealed the conversion of conjugates to single T cells and BSLB with evidence of T cell receptor (TCR) downregulation on T cells and TCR acquisition by BSLB as a function of α-CD3ε Fab density (Fig. 1B and Fig. S1A), as previously reported[5]. While BSLB acquired little CD2 and CD4, T cells acquired little fluorescent lipids (DOPE) indicating limited trogocytosis or trans-endocytosis of material from BSLB (Figs. 1B and S1B), contrasting to with has previously been reported for B cell IS [16].

**Figure 1.**
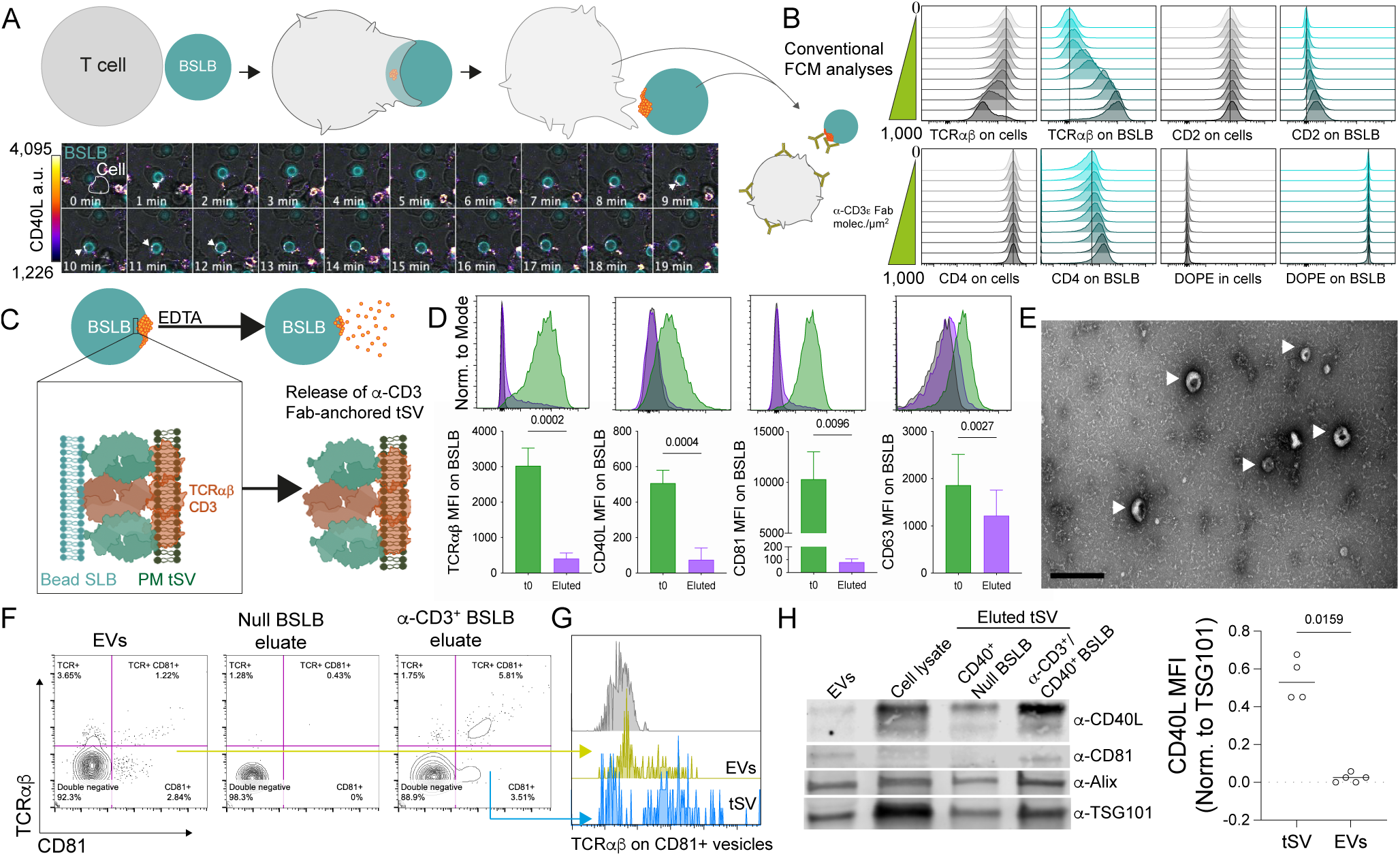
As synthetic APC, BSLBs trigger synapse formation and the release of tSV by stimulated T cells. (A) Representative time-lapse confocal microscopy showing the interaction between TH and BSLB and the active transfer of CD40L^+^ synaptic stamps to BSLB (white arrowheads), which are left behind after the interaction resolves. CD40L was tracked using 1 µg/mL of anti-CD40L clone 24-31 AF647. (B) FCM analyses of vesicular transfer to BSLB (teal histograms) from TH (grey histograms). BSLB were reconstituted with increasing densities of α-CD3ε Fab and 200 molec./µm^2^ of ICAM1 and 20 molec./µm^2^ of CD40. Vertical lines indicates the mode of cells and BSLB in the condition with no α-CD3 Fab (null). (C) After conjugate separation by cold, tSV are released from BSLBs with the use of 50 mM EDTA. (D) Elution results in the release of TCR^+^, CD40L^+,^ and CD81^+^, and to a lesser extent CD63^+^, from the surface of BSLB as measured by FCM staining comparing null BSLB (black histograms), with BSLBs prior to (green), and after elution (violet). (E) Transmitted electron microscopy of eluates reveals the presence of tSV (white arrows) and small soluble proteins. Scale bar= 200 nm. (F) Representative NanoFCM analyses showing the detection of TCRαβ^+^ and CD81^+^ events in isolated EVs (left panel), and the eluted material from Null BSLB (middle panel) and α-CD3^+^ BSLB (tSV, right panel). (G) overlaid histograms showing TCRαβ expression on CD81^+^ vesicles from EVs (yellow) and tSV (light blue) compared to double negatives (grey histogram). (H) Left: Immunoblot for comparison of CD40L, CD81, Alix and TSG101 in the EVs or eluted tSV deriving from 5 × 10^6^ of either cells or eluted BSLB, respectively. BSLB were previously cocultured with TH cells at 1:1 ratio. Whole cell lysate represents a total of 2.25 × 10^5^ cells. Right: CD40L levels in eluted fractions (tSV) and steadily released EVs as measured by immunoblot. Shown is CD40L corrected MFI (cMFI) normalised to TSG101 cMFI. Normality was determined using Shapiro-Wilk test and statistical significance was determined by two-tailed unpaired t-test (D) and by unpaired, two-tailed Mann-Whitney test (H). *p≤0.05, **p≤0.005, ***p≤0.0005, ****p≤0.0001 and ns = not significant. Data represents mean ± SEM of n=2 (A,F,G) and n=4 independent experiments (B-E, H).

We next eluted the tSV with ice-cold EDTA from BSLB previously enriched by low speed centrifugation (100*g* 1 min) and used differential and ultracentrifugation to concentrate the eluted tSV (see Methods). We focused our analyses on tSV containing the T cell effector molecules TCR and CD40L, the broad EV and PM marker CD81^+^ and on CD63^+^ tSV belonging to exosomes. Most of the TCR, CD40L and CD81 present on BSLBs was eluted under these conditions (Fig. 1C and D). Elution of CD63^+^ vesicles was less efficient but still significant (Fig. 1D). Electron microscopy images of eluted tSV revealed classical EV-like profiles (Fig. 1E, arrows). We were now able to compare tSV with steadily released EVs isolated from activated TH cells of the same donors cultured in EV-free media. TSV were larger than EVs by nanoparticle tracking analyses (NTA) (Fig. S1C). This was confirmed by nanoFCM revealing tSV having a median diameter of 82.13 ± 0.75 nm to 84.4 ± 5.99 nm, with or without CD40 in BSLB, respectively, compared to EVs from T cells of the same donors with a median diameter of 65 ± 25 nm. More detailed analyses using silica bead standards for size segmentation revealed a higher frequency of events larger than 113 and 155 nm in tSV compared with EVs (Fig. S1D). EVs showed a higher frequency of events in the 68 nm size bin (Fig. S1D). Comparison of marker-specific vesicles within tSV and EVs revealed that CD81^+^ tSV coexisted with TCRαβ^+^ at higher frequencies in tSV than EVs (Fig. 1F) and a larger median size for TCR^+^ tSV than TCR^+^ EVs (Fig. S1E,F), respectively. We also observed a larger median size for CD40L^+^ or CD81^+^ tSV than CD40L^+^ or CD81^+^ EVs (Fig. S1E), whereas no significant size differences were observed for BST2^+^ or CD63^+^ tSV and EVs. TSV also expressed higher levels of TCR or CD40L than EVs (Fig. 1G,H, and Fig. S1G,H). Since CD40L detection by antibodies is partly impaired by the presence of CD40 (see also Fig. 3B), we confirmed CD40L elution and compared its expression levels in tSV and EVs by immunoblotting (Fig. 1H and S1H). Most likely, the overall larger size and immune receptor content of tSV derives from the subpopulation of SE, which contains high densities of TCR and CD40L as microclusters >80 nm[5]. The membrane-anchored full-length CD40L was predominantly found (29.2 kDa) in the eluates of BSLBs, with neligible detection of its soluble ectodomain (Fig. S1H). Null CD40^+^ BSLB capture neligible amounts of CD40L from TCR-unstimulated cells, which might relate to the CD40-dependent capture of a fraction of EVs released in response to adhesion (i.e., ICAM1:LFA-1 interaction). The imaging of tSV markers with TIRFM and confocal imaging further revealed that beyond differences in size and immune receptor content, tSV displayed different degrees of co-localisation in intact, non-permeabilised ISs (Fig. S2A-I) reflecting their different subcellular origin and the compositional and spatial heterogeneity of tSV in the synaptic cleft.

TSV are likely composed of TCRαβ^+^, CD40L^+^ SE and CD63^+^ PE, which may have different release characteristics. We used the BSLB system to investigate requirements for tSV release using a panel of pharmacological inhibitors of candidate steps. A panel of 12 inhibitors targeting cytoskeleton dynamics and the transport of vesicular cargo from different organelles, such as multivesicular bodies (MVB) and the *trans*-Golgi network (TGN) was used (Table S1 and Fig. S3). Folling the treatment for 90 min at 37°C and 5% CO_2_, we monitored TCR, CD40L, BST2, and CD63 transfer from T cells to BSLB (Fig. S3A) and normalised trans-synaptic transfer to the maximum observed in untreated controls (%T_max_).

Actin cytoskeleton disruptors Latrunculin A and Jasplakinolide, but not Cytochalasin D, impaired cell-BSLB conjugates (Fig. S3B). Cytochalasin D and the other inhibirors didn’t impair cell-BSLB conjugate formation and thus allowed interrogation of distinct mechanistic steps in delivery of tSV cargo. Fortuitously, we observed a differential inhibition in the transfer of TCR^+^, CD40L^+^, BST2^+^ and CD63^+^ tSV from TH cells to BSLB, suggesting distinct mechansims of cargo delivery. The transfer of TCR^+^ tSV was selectively promoted by dynamin inhibition by Dynasore, and conversely, strongly reduced by inhibition of actin polymerization, ESCRT machinery, and neutral sphingomyelinases (Fig. S3C and Table S1). Reduced endocytosis or increased ubiquitination of TCR following Dynasore treatment [17] might contribute to the observed increase in TCR^+^ tSV transfer. The transfer of CD40L^+^ tSV was strongly reduced following acute inhibition of several machineries in the hierarchy TGN > actin dynamics and ESCRT > ceramide synthesis > dynamin > vacuolar H^+^-ATPases, and class I and II PI3K (Fig S3D). Other members of the TNF superfamily, such as FasL, have also been reported to be affected by interference with TGN trafficking [18, 19], suggesting a conserved mechanism of TNFSF delivery to tSV. In contrast, the transfer of BST2^+^ and CD63^+^ tSV was affected by inhibiting endosomal and lysosomal transport but not by inhibiting TGN transport (Fig. S3E-F). CD40L > TCR > BST2 showed a higher sensitivity than CD63^+^ tSV to the inhibition provided by the MG132-induced depletion of free-ubiquitin [5, 20, 21], suggesting that MVB-associated stores are unlikely to be affected by the acute pharmacological inhibition of the ESCRT machinery. The significant reduction of CD40L^+^ and CD63^+^ tSV transfer following N-SMases inhibition likely results from interference with broader membrane trafficking events involving both the TGN and endosomes where N-SMases preferentially locate to regulate membrane curvature and budding [22, 23]. Finally, our data suggest that compared to other vesicle subpopulations, the production of CD40L^+^ tSV requires the stepwise transport of CD40L to the PM and budding SE (Fig. S3G,H,J). While BFA and manumycin inhibited the α-CD3-induced upregulation of CD40L, CytD impaired CD40L^+^ tSV release despite its effective cell surface upregulation (Fig. S3D and S3H).

### T cell subsets have distinct tSV cargo

Next, we sought to evaluate whether different human T cells, including CD8^+^ cytotoxic T cells (CTL), CD4^+^ T helper cells (TH) and cultures enriched in CD127^low^CD25^high^ FoxP3^+^ cells (Treg), displayed distinctive TV transfer hallmarks. All cells were isolated from peripheral blood, activated, and further expanded *ex vivo* as detailed in Methods (see Fig. S4A-C). We evaluated the transfer of the TCRαβ heterodimer, CD2, the coreceptors CD4 or CD8, CD28, CD45, the tSV proteins CD63, CD81 and BST2, and effectors including regulatory enzymes (CD38, CD39 and CD73), and helper (CD40L) and cytotoxic (Perforin) proteins. Analyses were focused on the geometric mean or median fluorescence intensities of the population of single T cells and single BSLB after gentle dissociation of conjugates (Fig. 2A). A normalised synaptic transfer metric was defined as the percent enrichment of T cell proteins on single BSLB from the total mean signal observed in single BSLB and single cells (Fig. 2A and ref.[5]).

**Figure 2.**
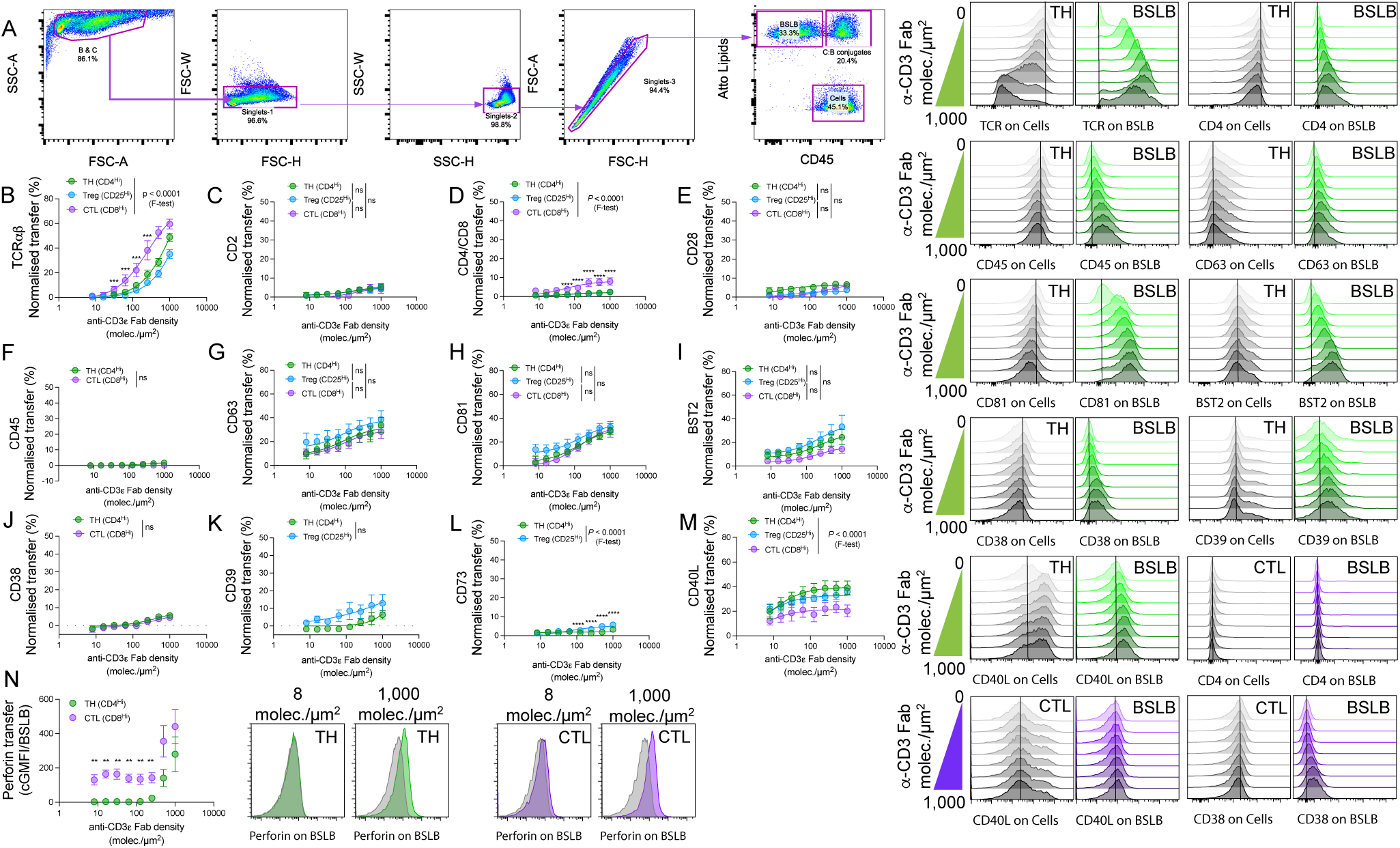
The synaptic transfer of vesicular effectors relates to the functional properties of different T cell subsets. (A) Gating strategy for selecting single BSLB and T cells to measure the transfer of vesicular effector at the IS. BSLBs presenting α-CD3ε Fab (0 to 1,000 molec./µm^2^), 200 molec./µm^2^ of ICAM1 and 20 molec./µm^2^ of CD40 were used in all experiments. (B-L) Percent Normalized Synaptic Transfer (NST%= [cGMFI_BSLB_/(cGMFI _BSLB_ + cGMFI_Cells_)]*100) to BSLB from TH (green), Treg (blue) and CTL (violet). Ligands included (B) the antigen receptor heterodimer TCRαβ; (C) CD2; (D) TCR co-receptors (CD4 or CD8); (E) CD28; (F) CD45; (G) CD63; (H) CD81; (I) BST2; (J) CD38, (K) CD39; (L) CD73, and (M) CD40L. (N) Perforin (PRF1) transfer as background corrected GMFI is shown (null BSLB-subtracted). Representative histograms for most makers are shown on the right for TH (green) and CTL (violet). Statistical significance was determined by multiple t-test comparing tSV transfer across different α-CD3ε Fab densities and among different T cell populations. α-CD3-Fab EC_50_ and the marker transfer maximum (T_max_) were determined using F-test and three to four parameters fitting (H-J). *p 0.05, **p≤ .005, ***p≤0.0005, ****p≤0.0001 and ns = not significant. Data represent means ± SEM of n=4-10 independent experiments.

Because we reconstituted our BSLBs in the absence of ligands for TCR coreceptors CD4/CD8 (i.e., pHLA) and CD2 (i.e., CD58), we therefore expected the vesicular shedding of TCR to include limited coreceptors and CD2 (Fig. 2B-D). Compared with TH, CTLs transferred significantly more TCR and coreceptor in the range of α-CD3ε-Fab densities tested and consistently showed smaller α-CD3ε-Fab EC_50_ values of 251.2 for TCR (compared with 1,251 in TH) and 50.87 for the coreceptor (compared with 56.44 in TH) (Fig. 2B-D). The higher antigen sensitivity of CTL compared with TH and increased physical proximity between TCR and CD8 [24], might promote their sorting in tSV. Similarly, using chimeric antigen-receptor expressing T cells (CART) specific for HLA-A*02: NY-ESO-1_157-165_ we tested whether the T1 CAR displays similar transfer dynamics than TCR. Interestingly, we observed a significantly higher transfer of the CAR than TCR at comparable densities of α-CD3ε-Fab and agonistic HLAp complexes (Fig. S4D). CART also transferred higher amounts of CD8, suggesting that CAR triggering by agonistic pHLA, even at low densities, promotes the efficient release of CAR and CD8 (Fig. S4E). Like CD2 and coreceptors, and despite their high level of cell surface expression, we observed a lack of CD28 and CD45 enrichment in the tSV released by different T cell subsets (Fig. 2C-F). While CD45 is normally excluded from the cSMAC and the synaptic cleft, the coreceptors, CD2 (at exceptionally low levels) and CD28 require binding to cognate ligands to be mobilised as microclusters to the synapse centre [25–27]. The latter suggests that tSV heterogeneity is partly defined by APC ligands dynamically feeding information to T cells, which we explored further in the next section. The limited enrichment of CD4, CD8, and CD45 in tSV makes them suitable secondary parameters for discriminating single BSLBs in FCM analyses and sorting (Figs. S1A, S4I, and S6A).

TH, CTL, and Treg transferred comparable levels of CD63^+^, CD81^+^, and BST2^+^ tSV to BSLB (Fig. 2G-I), indicating conserved mechanisms delivering these components to the synaptic cleft. Consistently, CART showed a comparable transfer of CD63^+^ tSV when stimulated through the CAR and the TCR (Fig. S4F). We next measured the vesicular transfer of effector ectoenzymes, namely the ADP-ribosyl cyclase CD38 and the AMP- and adenosine-producing ectonucleotidases CD39 and CD73 (Fig. 2J-L). While CD38 polarises to the IS[28] and participates in the generation of T-dependent humoral immunity[29], CD39 and CD73 are known tolerogenic mediators [30–32]. TH and CTL showed a limited, conserved, and comparable transfer of CD38^+^ tSV to BSLB (Fig. 2J). Treg, on the other hand, showed an increased transfer of CD39 (T_max_ = 17.83% compared with 6.5% in TH; Fig. 2K), and CD73 (T_max_ = 5.73 % compared with 1.9% in TH; Fig. 2L), which associated with a higher central clustering of CD39 and CD73 in the cleft of Treg synapses (Fig. S4J-K). The inclusion of itinerant enzymes such as CD38, CD39, and CD73 in tSV, although limited, might exert a feed-forward regulatory effect as shown for tumor-derived sEV[33].

We also compared the transfer dynamics of CD40L and perforin, two major effectors mediating the activation and killing of APCs, respectively. Upon TCR triggering, we observed a conserved relative transfer of CD40L in T cells, with TH secreting the most (Fig. 2M, see also Fig. 3). Conversely, compared to TH, CTLs showed a reduced threshold for perforin release, which was transferred to BSLB at α-CD3ε-Fab densities as low as 8 molec./µm^2^ for CD8^+^ T cells (Fig. 2N). In CART cells, perforin release followed a different behaviour than exocytic CD63^+^ tSV. More specifically, TCR-elicited perforin deposition on BSLB was significantly higher than that resulting from CAR-triggering (Fig. S5D), suggesting differential mechanisms influencing the release of tSV and cytotoxic SMAPs at the IS. Altogether, our data indicate that 1) T cell subsets display differential dynamics of tSV transfer consistent with their effector functions and receptor-ligand interactions occurring at the IS and that 2) BSLB are excellent tools to study the biogenesis of both tSV and perforin-containing SMAPs.

**Figure 3.**
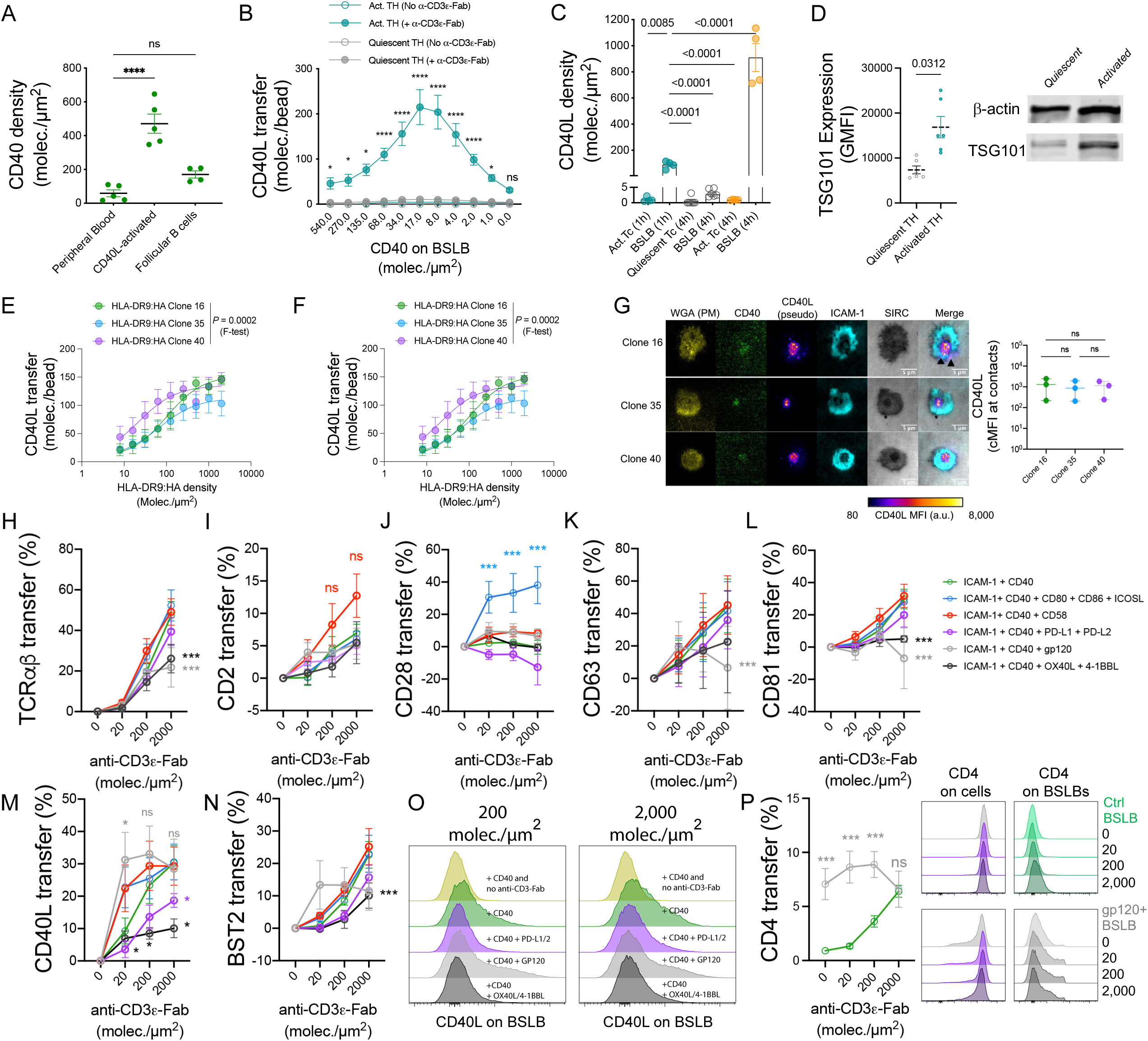
BSLBs reveal the dynamics influencing the biogenesis of CD40L^+^ tSV. (A) CD40 densities on the surface of human B cells isolated from either peripheral blood (CD19^hi^ HLA-DR^+^ and CD40^+^) or palatine tonsils (also CXCR5^hi^). (B) Based on data in A, BSLBs were reconstituted with increasing densities of human CD40 and analysed for their capacity to instigate the release of CD40L^+^ tSV. Quiescent and α-CD3/CD28-activated TH are compared. BSLBs presenting either no α-CD3ε or 1,000 molec./µm^2^ were used in these experiments. (C) As in B, TH cells were incubated for 4h with BSLB and the density of CD40L compared to 1 h activated TH controls. (D) TSG101 expression by flow cytometry (left) and immunoblotting in quiescent and activated TH. (E and F) Comparison of CD40L^+^ tSV transfer among different T cell types (*green:* TH; *blue:* Treg-enriched; *violet:* CTL), and among T cell clones (E) responding to the influenza H3 HA_338-355_ peptide presented in HLA-DRB1*09:01. (G) TIRFM images showing CD40L clustering within the synapses of T cell clones stimulated on SLB containing 30 molec./µm^2^ of antigen, 200 molec./µm^2^ of ICAM1 and 20 molec./µm^2^ of CD40. Scale bar = 5 µm (A-H) After cell: BSLB conjugate dissociation FCM measurement was performed on BSLB to calculate the NST% of tSV positive for (H) TCR, (I) CD2, (J) CD28, (K) CD63, (L) CD81, (M) CD40L and (N) BST2 was measured on control (green) or BSLB presenting CD80, CD86, ICOSL (blue), CD58 (red), PD-L1 and PD-L2 (magenta), HIV-1 gp120 (grey), and OX40L and 4-1BBL (black) (all at 100 molec./µm^2^). (O) Representative half-overlaid histograms showing CD40L^+^ tSV transfer to ligand- and α-CD3ε-Fab-coated BSLB. Control BSLB were tailored to present ICAM1, CD40 and titrations of α-CD3ε-Fab. (P) NST% of CD4 to control (green) and HIV-1 gp120-presenting BSLBs (grey; red arrows indicate the loss of CD4 on TH). Normality was determined using Shapiro-Wilk test, and statistical significance was determined by multiple t-test to compare levels of CD40L^+^ tSV transferred to BSLB across different CD40 densities, times, T cell populations (B-C, E-F), and increasing antigen densities (either α-CD3ε Fab or HLA-DRB1*09:01:HA complexes). α-CD3ε-Fab EC_50_ and marker maximum transfer (T_max_) were calculated using three to four parameters F-test (E, F). In D, Wilcoxon matched pairs signed rank test. *P* = 0.0312 was used to compare TSG101 expression between quiescent and activated cells. In H-P, multiple t-test comparing control BSLB presenting 200 molec./µm^2^ of ICAM1, 20 molec./µm^2^ of CD40 and α-CD3ε Fab (0 to 2,000 molec./µm^2^) with BSLBs also presenting the selected group of proteins each at 100 molec./µm^2^. *p≤0.05, **p≤0.005, ***p≤0.0005, ****p≤0.0001 and ns = not significant. Data represents means ± SEM of n=4 (A-E and H-P) and n=3 (F, G) independent experiments.

### IS composition regulates SV cargo

In the dynamic exchange of information between T cells and APCs several factors might influence the effector cargo of tSV. Hence, we harnessed BSLB to study the dynamics influencing CD40L^+^ tSV release under near-physiological conditions and using absolute quantifications (please refer to Methods). Absolute measurements offer a better comparison of varying CD40L expression levels among different T cell subsets (e.g., Fig. 2M). First, we measured CD40L^+^ tSV release in response to CD40 densities of different primary human B cell subsets (minimum and maximum values of 16 and 646 molec./µm^2^; Fig. 3A). BSLB covering a range of 0 to 540 molec./µm^2^ of CD40 were used to compare the tSV transfer dynamics to BSLB between previously activated (and expanded) TH blasts and freshly isolated (quiescent) TH. As shown in Fig. 3B, while quiescent TH transferred negligible amounts of CD40L, activated TH showed high sensitivity to CD40, transferring CD40L to CD40 densities as low as 2 molec./µm^2^. However, this high efficiency reached a maximum transfer (T_max_) at around 20 molec./µm^2^ of CD40, which at higher densities outcompeted the anti-CD40L detection antibody [5]. The latter is expected as CD40 displays a high affinity for its ligand (0.5-7.13 nM [34]). Therefore, we used the physiological minimum of CD40 (20 molec. /µm^2^) to study the dynamics affecting CD40L sorting in tSV. We used a vesicle stamp area of 0.84 µm^2^ [5] to estimate and compare the effective densities of CD40L on the vesicular stamps of BSLB. Even after 4h, no significant transfer of vesicular CD40L was observed from quiescent cells (Fig. 3C). In contrast, activated TH blasts transferred significant amounts of CD40L and led to densities up to 1,000 times those found on the PM, suggesting a significant gain in CD40L binding valency resulting from its vesicular packing. Remarkably, higher CD40L^+^ tSV transfer in activated cells related to their higher expression of TSG101 (Fig. 3D), indicating that T cell activation leads to the simultaneous up-regulation of effectors and ESCRT-I. Further comparison of TH, Treg, and CTL revealed phenotype-specific differences in the dynamics of CD40L transfer among activated T cells, with the amounts released in the order TH>Treg>CTL (Fig. 3E). As in Fig. 2L, TH showed increased sensitivity for CD40L release compared with Treg and CTL (F-test, p<0.0001).

Next, we addressed whether TCR:pMHC binding potency influences the release of CD40L^+^ tSV. We used 3 TH clones specific for the same influenza H3 hemagglutinin peptide HA_338-355_ (NVPEKQTRGIFGAIAGFI) but displaying different TCR:pMHC potencies as determined by an IFN-γ release assay (Table S2). T cell clones showed comparable levels of CD40L transfer at higher antigenic HLA densities and formed morphologically similar synapses (Fig. 3D-E). However, contrary to their IFN-γ response, clones showed differences in the EC_50_ for CD40L transfer with clone 40 displaying the lowest (*P*=0.0002, EC_50_= 22.02, compared to 66.01 and 101.9 for clones 16 and 35, respectively; Table S2), suggesting that TCR: pMHC affinity has more substantial effects on the release of CD40L^+^ tSV at lower antigen densities. The latter phenomenon might underpin the maintenance of an active CD40L-CD40 pathway in autoimmune diseases, where high densities of self-antigens might suffice to trigger the release of CD40L^+^ tSV despite low affinities of TCR for self-peptides and the assembly of non-canonical immune synapses [35].

Beyond antigens (signal one), APCs also present membrane-bound accessory signals (signal two) to fine-tune the activation of engaged T cells. Because the influence of accessory molecules on the vesicular output of the synapse is not fully understood, we next sought to evaluate the effects of different known T cell costimulatory and repressing signals on the shedding of CD40L^+^ and other tSV. We incorporated costimulatory signals including CD80, CD86, ICOSL, and CD58, the latter of which forms a distal domain boosting TCR signalling known as CD2 corolla [27]. We also included the negative regulators PD-L1 and PD-L2 and other TNF ligand superfamily (TNFSF) members, including OX40L (TNFSF4) and 4-1BBL (TNFSF9), whose interplay with the CD40-CD40L dyad remains poorly characterised. To evaluate whether the synaptic output of T cells could also be affected by foreign proteins, such as those participating in virological synapses, we included the HIV-1 gp120 protein, which is a known high-affinity ligand for CD4 [26, 36–38]. We then compared the differences in NST% between different BSLB compositions using those containing only ICAM1, CD40, and a titration of α-CD3ε-Fab as controls (Fig. 3H-P). While CD58 incorporation slightly promoted transfer of CD2 (Fig. 3I), incorporation of CD80, CD86 and ICOSL induced an increased synaptic transfer of CD28 (Fig. 3J), and in a lesser extent that of CD40L (Fig. 3M). Most remarkably, the inclusion of PD-L1 and PD-L2 on BSLB resulted in a significant reduction in the transfer of CD40L^+^ tSV (Fig. 3M), and a less significant reduction of TCR^+^, CD28^+^, CD63^+^, CD81^+^, and BST2^+^ tSV. Similarly, BSLB loaded with a combination of 4-1BBL and OX40L impaired the transfer of vesicular TCR, CD81, CD40L, and BST2 from TH (Fig. 3H, L-N). Whether these differences relate with the competitive occupancy of intracellular signalling adaptors shared between CD40L, OX40 and 4-1BB are subject of further investigation. BSLB loaded with the HIV-1 gp120 instigated a reduced transfer of TCR^+^, CD63^+^ and CD81^+^ tSV at high α-CD3ε Fab without significantly affecting the transfer of CD40L^+^ tSV (Fig. 3H, K-M). Importantly, gp120 promoted CD4 transfer to BSLB (Fig. 3P) even without TCR triggering. As previously shown, CD4 transfer might be supported by TCR-like signalling events involving Src, Lck, and CD3ζ phosphorylation [26]. The gp120-induced transfer of CD4 and the CD80/86-instigated transfer of CD28 provide further evidence linking ligand binding with the sorting of proteins in tSV. Altogether these data provide proof of concept that as synthetic APCs, BSLBs facilitate the study of cognate and non-native molecular interactions occurring at the cell-bead interface and shaping the composition of T cell tSV.

Next, we used CRISPR/Cas9 genome editing to test the functional relevance of endogenous elements participating in tSV biogenesis and the inclusion of CD40L. Because we used anti-CD3 Fab for the HLA-independent triggering of TCR, we expected negligible participation of CD4 in tSV biogenesis. Hence, we selected a guide RNA (gRNA) producing downregulation of the coreceptor (67.83 ± 10% of baseline) as a control. We also selected gRNAs targeting membrane structural proteins (*BST2*, *CD81*) and the ESCRT-I component *TSG101*, which participates in the budding of SE [5]. We also targeted ADAM10, a disintegrin and metalloproteinase mediating the shedding of surface FasL and CD40L from activated T cells [39, 40], as it might mediate CD40-induced transfer of soluble CD40L trimers [41]. Knockout of the relevant targets was observed with some residual surface on cells edited for BST2 (45.08 ± 23.19% of control), CD81 (7.63 ± 4.8% of control), and ADAM10 (9.37 ± 2.83% of control). ADAM10 downregulation led to a significant increase of cell surface CD40L (224.2 ± 114.8%) (Fig. S5A-D). We compared tSV transfer in terms of the T_max_ % observed in controls (*CD4*-gRNA). While ADAM10 downregulation increased both the cell surface expression and the transfer of CD40L^+^ tSV to BSLB, the downregulation of CD81, BST2 and TSG101 reduced the transfer of CD40L^+^ tSV without altering the baseline expression levels of CD40L at the PM (Fig. 4A and S5A-B). As expected, *CD81* and *BST2*-edited cells showed significant reduction in the transfer of their respective CD81^+^ and BST2^+^ tSV (Fig. 4D-E, respectively), and non-significant differences with regards to other markers (Fig. 4B-E). *TSG101* editing led to a significant reduction of CD40L^+^ and TCR^+^ tSV transfer without affecting their baseline surface expression levels, indicating a bona-fide inhibition of their vesicular sorting dependent on ESCRT-I (Fig. 4A-B). TSG101 downregulation partly phenocopied the effects of ubiquitin depletion by MG132, with inhibition of transfer affecting mostly CD40L^+^ tSV, and to a lesser extent TCR^+^ tSV (Fig. 2C-D). CD63^+^ tSV transfer was less affected by downregulation of CD81, ADAM10, BST2 and TSG101 (Fig. 4C), providing an additional piece of evidence for CD63 vesicular transfer relating mostly to the transport and exocytosis of preformed MVB, which is also congruent with our observations with the panel of inhibitors. We next sought to evaluate whether reduced tSV transfer to BSLB relates with impaired trafficking and central clustering of CD40L at the T cell-SLB interface. We compared CD40L total FI within synapses (i.e., as median IntDen per donor) as a function of the Tmax of controls. Imaging of ISs formed on planar SLB further revealed that reduced transfer to BSLB correlated with a significant reduced total clustering of CD40L within synapses of *CD81-* and *TSG101*-edited cells, with a less stronger effect observed in *BST2*-edited cells (Fig. 4F-G). When analysing the dynamics of tSV transfer to BSLB we observed a reduced T_max_ of CD40L^+^ tSV for *CD81-*, *BST2-*, and *TSG101-*edited cells, the latter of which also displayed a reduced T_max_ for TCR^+^ tSV (Fig. S5F-I). *TSG101*-edited cells also showed reduced upregulation of cell surface CD40L upon TCR-triggering (Fig. S5J), consistent with the reduced total clustering of CD40L at the IS interface. Expectedly, *ADAM10*-edited cells showed increased T_max_ and reduced α-CD3ε-Fab EC_50_ for the release of CD40L^+^ tSV, and a marked cell surface upregulation of CD40L following α-CD3ε stimulation (Fig. S5I and J, respectively). The partial reduction in CD40L transfer following downregulation of CD81 and BST2, suggests that rather than being essential, these integral membrane proteins work as aiding factors in the release of CD40L^+^, but not TCR^+^ nor CD63^+^ tSV. Also, the differential requirements for TSG101 among TCR^+^, CD40L^+^ and CD63^+^ tSV, which is phenocopied by MG132 treatment, demonstrates different ESCRT-I requirements among tSV. The increased transfer of CD40L^+^ tSV from ADAM10-deficient cells also suggests minimal inclusion of cleaved ectodomains in the transferred pool of CD40L, which is also supported by our immunoblot analyses (Fig. S1G-H).

**Figure 4.**
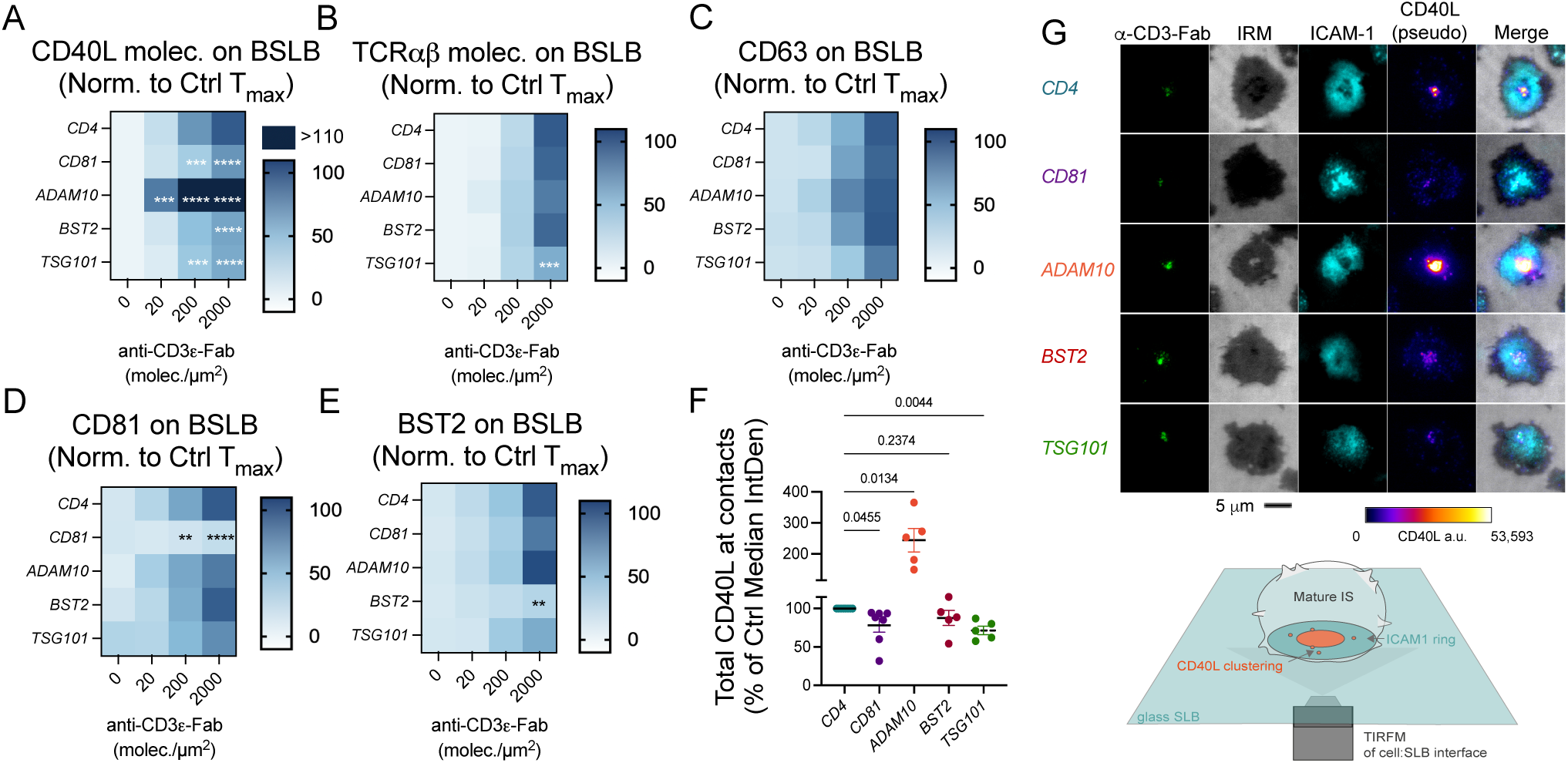
Harnessing BSLBs to identify endogenous proteins participating in the biogenesis of CD40L^+^ tSV. (A-E) Data showing the transfer of tSV as measured by flow cytometry of BSLB after the dissociation of cell:BSLB conjugates and expressed as percent of the maximum transfer (T_max_%) observed in *CD4*-edited controls. Heatmaps show T_max_% for different tSV populations, including those positive for (A) CD40L, (B) TCRαβ, (C) CD63, (D) CD81, and (E) BST2 on BSLB incubated with different CRISPR/Cas9-edited cells (targets shown in italic). BSLB contained 200 molec./µm^2^ of ICAM-1, 20 molec./µm^2^ of CD40 and four increasing densities of anti-CD3ε Fab (0-2,000 molec./µm^2^). Normality was determined using Shapiro-Wilk test and statistical significance was determined by multiple t-test corrected for multiple comparisons (Holm-Sidak) and comparing tSV transfer to BSLB between CRISPR/Cas9 edited cells and *CD4*-edited controls (depicted as % of CD4-edited T_max_). *p≤0.05, **p≤0.005, ***p≤0.0005, ****p≤0.0001 and ns = not significant. (F-G) TIRFM imaging and quantification of total CD40L FI in the synapses formed by CRISPR/Cas9 genome-edited cells. (F) Median CD40L integrated density (i.e. total FI) in immunological synapses of CRISPR/Cas9-edited cells as % of the maximum observed. Each circle represents a donor. Means of medians were compared using Mixed-effect analysis with geisser-greenhouse correction and Fisher’s LSD test. (G) Top: Representative TIRFM images showing mature immunological synapses and the levels of centrally clustered CD40L. Bottom: diagram depicting the imaging of CD40L and other interfacial signals in IS formed between T cells and stimulating SLB. Data representative of n=6 (A-E) and 5 (F-G) independent experiments.

### tSV carry distinct miRNAs from EVs

Within the framework of harnessing BSLB to study particulate effectors released at the IS, we then sought to identify miR species associated with tSV. To minimize cell contamination, we isolated BSLB coated with TCR^+^ tSV using fluorescence-activated cell sorting of single beads after 90 min of culture with T cells (Fig. S6A), followed by small RNA purification. Small RNAs were also purified from EVs from the same donors as controls. Prior to comparing the RNA species identified in cells, EVs and tSV, we used the less abundant RNA species found in null BSLBs (Fig. S6A, grey overlaid histogram) as background for the identification of miR enriched in tSV. Analyses of tSV and EVs libraries’ raw reads revealed a similar composition of small RNA species, of which miRs were overall more enriched in tSV than EVs and the parental cells (Fig. 5A). After identifying small RNA species conserved across samples, donors, and sequencing runs, GSEA further revealed public and tSV- or EVs-specific miRs in both TH and CTL. Of 200 miR species enriched in TH, 11 were uniquely enriched in tSV and 23 in EVs, whereas of 163 miR species enriched in CTL, 17 were unique to tSV and 19 to EVs (Fig. 5B and Table S3). Consistent with the wider variety of enriched miR in sEV, we observed a higher variance compared to tSV (Fig. 5C). Analyses of enriched motifs with MEME [42] identified motifs to unique and shared miR species in TH (Fig. 5D) and CTL (Fig. 5E), which together redundantly associated with GO pathways related to immune cell signalling and anti-tumor immunity (Fig. 5F and heatmaps in Fig. S6B). Congruently, miR target enrichment analyses of enriched miR further revealed a similar degree of redundancy in the identified targets (see Table S4 for the full list of targets and Fig. S6C-H for GO terms), which included RNA-binding proteins, such as AGO1/2, DICER, GRB2, GRAP2, PLAGL2, RRM2, TNRC6A/B, and YBX1; immune receptor adaptors and signalling proteins such as CD2AP, IFNARs, SMADs; and synaptic ligands and effectors including ICAM1 and thrombospondin-1 (THBS1), among others.

**Figure 5:**
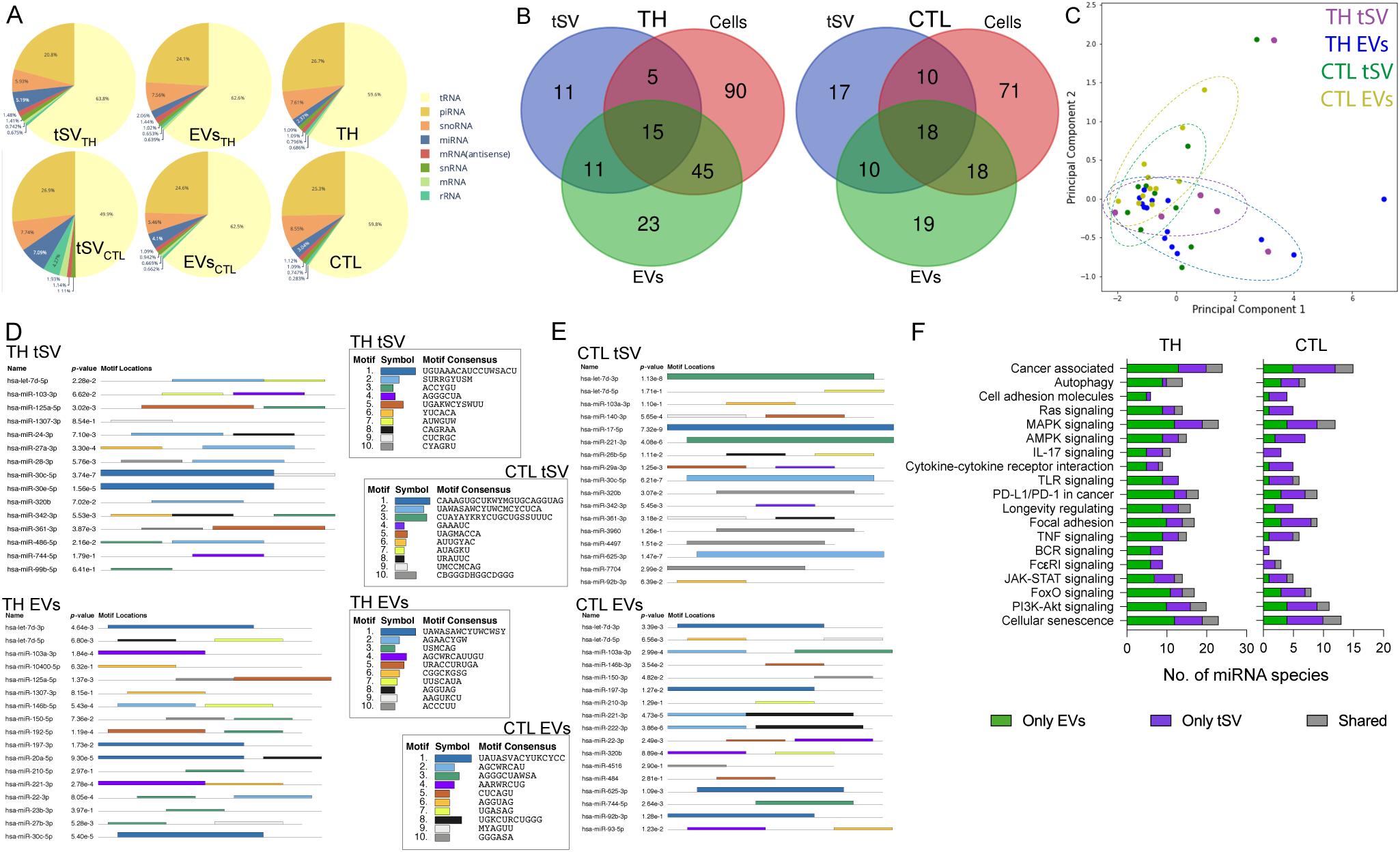
tSV are enriched in miRs, including both private and public species with considerable functional redundancy to EVs. (A) Abundance of different genome-mapped RNA species found associated with tSV, syngeneic steadily released EVs, and parental cells. (B) Venn diagram showing the overlap of enriched miR species associated with tSV, EVs and parental cells (see also Table S3). (C) miR heterogeneity is further reflected by PCA of the analysed tSV and EVs. (D-E) Motif analyses using MEME identified consensus sequences (central inserts) mapped across different miRs in TH tSV (D, top), TH EVs (D, bottom), CTL tSV (E, top) and CTL EVs (E, bottom). (F) Bar graph displaying the number of miR species enriched uniquely in tSV (violet), uniquely in EVs (green), or shared (grey). TH miR targets are shown in the top panel, and CTL miR targets are shown in the bottom panel. Data representative of at least six independent experiments, samples with low RNA yields were pooled for sequencing (n = 8 donors).

Although EVs showed slightly more miR species, the overall percent enrichment of these small RNA species was higher in tSV. We next sought to evaluate whether the increased percent enrichment of miR in tSV relates to a higher RNA binding protein (RBP) content in these vesicles. Following protein capture, digestion and elution using S-Trap, we performed label-free quantifications with liquid chromatography-tandem mass spectrometry (LC-MS/MS) on tSV and EVs isolated by differential centrifugation of BSLB eluates and cell-cleared culture supernatants, respectively. Since contaminants derived from the elution reagents and procedure might impact the quality of our comparisons, we first identified proteins enriched in eluates derived from activating (α-CD3 Fab^+^) BSLB comparing to null BSLB (coated only with ICAM1 and CD40)(Fig. S7A). With the aid of Pegasus we generated a list of differentially expressed proteins to compare with the EVs from the same TH and CTL cells, respectively (Fig. S7B, C). PANTHER based GSEA revealed vesicle-specific differences between eluates (tSV) and EVs, including the reduced detection of ADAM10 and contaminants Histone 3, LGALS3BP [43], and RNAse 4 in tSV (Fig. S7D), and the high enrichment of targets predicted to interact with tSV miRs, such as SMAD4, DNMT1 and GRB2 in tSV (Fig. S7E and Table S4). Proteins related to global nucleic acid-binding, including RNA-binding and RNA-interference, stood out in tSV compared with EVs (Fig. S7F, G), with an overall difference in the enriched proteins found in tSV and EVs. Among them, YBX1, a known protein packing small noncoding RNAs (sncRNA) in EVs [44, 45] was identified as a common RNA-binding protein enriched in tSVs (TH>CTL; Fig. S7F), as well as with > 3-fold enrichment in SE tethered to sorted BSLBs (see ref. [5]). Other RNA-binding proteins were also commonly identified in independent LC-MS/MS runs of eluted tSV and subpopulations of SE anchored to sorted BSLB (Fig. S7H). To validate these findings, we then performed TIRFM imaging of TH cell synapses to localise the distribution of some of these RBPs. The localization of in YBX1 and, in a lesser extent, the SE-enriched SF3B3[5] in both the synaptic cleft and the proximal membrane of T cells interacting with α-CD3ε further validated our LC-MS/MS findings (Fig. S7I and J, respectively, compare with isotype control in K). YBX1 localised as discrete puncta both in the synaptic cleft and in vesicular tracks left behind by cells spontaneously breaking synapse symmetry and resuming migration (compare with 7I, white arrows). Supporting the active release of vesicular RNA, we further found the polarisation of ER-like structures and release of ceramide^+^ and RNA^+^ puncta in both the synaptic cleft and in vesicular tracks of live, non-permeabilised TH IS (Fig. S7L,M, yellow arrows). As evidenced by the super-resolution imaging with eTIRF-SIM, most RNA puncta lined the periphery of the synaptic cleft as early as 5 minutes during synapse assembly, indicating the rapid mobilisation of RNA^+^ compartments to the contacting membrane of stimulated T cells (Fig. S7M, right panel). The second layer of information provided by transferred miR and their protein partners might be pivotal for the functional modulation of APCs and the fine tuning of T cell activation and effector function by serving as a mechanism for suppressor miR elimination. Altogether, the polarisation and central clustering of immune receptors, ER-like structures, RNA and YBX1 to the synaptic cleft enriched in trans-synaptic effectors supports the superior role of the IS in the transfer of a broad range of intercellular messengers.

## DISCUSSION

The first observation of T cells being successfully activated by bead-supported bilayers presenting antigenic MHC-II dates back to 1986 [46]. Others have demonstrated the versatility of BSLB as APCs in several experimental contexts, including measuring the dimensions of interfacial receptor-ligand interactions and the localisation of lipid species and CD3ε in the interacting pole of T cells [47, 48]. Here we leveraged the APC capabilities of BSLB to isolate and study in more detail the composition and biogenesis of T-cell released EVs. Most importantly, BSLB facilitated novel comparisons of tSV and EVs, providing evidence for highly heterogeneous yet unique populations of vesicles being released in the limited dwell-time of the IS. We corroborated quantitative differences in size, protein loads and association to miR and RBPs between tSV and EVs. Also, the rate of secretion is higher for tSV compared with EVs. We have previously shown that TH IS shed an approximate of 25-30 TCRαβ^+^ SE after 20 min of IS formation on SLB[4, 5]. In contrast, the quantification of TCRαβ^+^ EVs by NanoFCM evidenced their rather low constitutive secretion ranging from a minimum of 0.54 to a maximum of 8.96 EVs/cell over a period of 48h of culture.

The acute pharmacological inhibition and CRISPR/Cas9-editing of TH strongly indicated that mechanistically tSV heterogeneity is partly explained by contributions of several cellular machineries recruited at the interacting pole of activated T cells. We provide evidence for an overarching hierarchy in the mechanisms participating in the vesicular transfer of CD40L, with dominant participation of ER/TGN transport and ESCRT machinery, and secondary involvement of MVB and lysosomes. The selective increase in TCR^+^ tSV transfer resulting from dynamin inhibition and the reduced effect of ESCRT-inhibition/downregulation on CD63^+^ tSV transfer provide additional evidence for different sorting mechanisms participating in the biogenesis of tSV (Figs. S3C-F and 5). We provide evidence for a critical role of ADAM10, a protease known to be active at the PM, in regulating the pool of surface CD40L and the resulting levels of CD40L^+^ tSV shedding dependent on CD40. Interestingly, our LC-MS/MS analyses indicate that compared to tSV, EVs incorporate a higher amount of ADAM10 (Fig. S7D), explaining their reduced vesicular CD40L (Fig. 1H).

Microvilli lead the formation of effector membranous particles referred to as synaptosomes enriched in LFA-1, CD2 and cytokines [49]. Microvilli tips accumulate proteins including TCR, CD2 and CD4 [49, 50]. Here we show that T cells transfer little vesicular CD2 and CD4 (Fig.1B and 2C,D), which increased at limited levels only upon ligation by CD58 and HIV-1 gp120, respectively (Fig. 3I and 3P). Also, TCR signalling, which is required for tSV biogenesis, is known to produce microvilli disassembly [50], suggesting that synaptosomes differ in origin and structure from ectosomes and contribute minimally to the vesicular output of the IS. Rather, synaptosome-like structures might mediate the negligible release of EV-like material to BSLB sustaining cell adhesion but devoid of antigen/α-CD3 Fab (Fig. 1H).

TSV biogenesis is a highly dynamic process connecting the integration of signals presented by APCs with the particulate output of activated T cells. Earlier electron microscopy studies demonstrated budding of TCR^+^ SE with partial polarisation of MVB and TGN in stimulated cells (figure 1C in ref[4]). In these experimental settings, SLB lacked CD40, which as we later demonstrated, promotes quick mobilisation of CD40L to the PM and into budding SE as microclusters [5]. Here we provide evidence that shedding of CD40L^+^ tSV requires CD40 on presenting BSLBs and its quick mobilisation from the ER-Golgi. We provide evidence of other signals finely tuning tSV heterogeneity, including B7 receptors (CD80/86), PD-1 ligands, other TNFSF members, and non-native ligands. For instance, HIV-1 gp120 promoted the vesicular transfer of CD4 and CD40L at low physiological densities of α-CD3ε Fab. Similarly, the non-native interaction between CARs and antigenic HLA complexes led to a remarkably efficient transfer of CAR as part of tSV and at the expense of its cell surface expression levels (Fig. S4D, red arrow). The superior vesicular shedding of CAR containing CD3ζ signalling tails might result from rewired TCR signalling networks and the formation of multiple short-lived synapses [51]. Since reduced CAR expression might desensitise CART to new encounters with target cells, more research is needed to understand the impact of CAR release on the therapeutic effectiveness of adoptive T cell therapies.

The small RNA sequencing of BSLBs made possible the first side-by-side comparison of the miR cargo associated with tSV and EVs. We found that both miRs and RBPs were significantly more enriched in tSV than EVs (Fig. 5A and S7F,G). The high level of target redundancy found in the enriched miRs of both tSV and EVs provides an alternative mechanism supporting the effector role of miRs, especially when few copies of single species are transferred associated with EVs [52]. In other words, unlike immune receptors, which require gains in avidity to feed-forward activation signals to APCs, miRs might rely on redundancy to modulate the function of APCs. We found RBPs enriched in LC-MS/MS of tSV and SE, namely YBX1 and SF3B3, in the synaptic cleft of TH, implying the transfer of miR ribonucleoprotein complexes across these short-lived cellular contacts. Since miR targets participate in cell cycle regulation, senescence, and immune receptor signalling, the release of tSV might also promote the clearance of miR species controlling various aspects of the T cell effector response. Interestingly, telomeric binding proteins were also found among the enriched GO terms of tSV (rank 27 in Fig. S7G) and the differential expression analyses of SE proteins [5]. Whether the transfer of miRs, and to a lesser extent telomeres, in tSV contributes to the paracrine regulation of senescence, as preliminarily shown for APCs transferring telomeres [53], remains to be elucidated.

How sncRNAs associate with tSV is not fully understood. The recent identification of N-glycosylated sncRNA species on the cell surface [54] suggests that if such modified species do exist, they might be transferred associated with the glycocalyx of synaptic ectosomes. Since we have observed the shedding of glycan-rich particles within the synaptic cleft ([5, 6] and WGA puncta in Figs. S2B-I, 3G, S4K), we are studying whether glycoRNAs are loaded on the surface of tSV.

The intrinsic feed-forward function of tSV-enriched immune receptors and miRs, specifically and timely released in response to antigen is different in kind and structure to other signals integrated during early T cell activation. Therefore, we consider that the heterogeneity and effector diversity of the particulate output of T-cell synapses, including tSV and SMAPs, deserves its functional classification as signal four. We expect BSLB will help others in the high throughput study of other key factors driving signal four-dependent communications. Undoubtedly, the comparison of EVs and tSVs in other immune cells known to secrete highly heterogeneous EVs, such as DCs [55], or in cells using similar interfacial communications (e.g., CD40L^+^ platelets and stromal cells secreting ADAM10 inhibitors) is needed to elucidate whether tSV are pivotal information units of cellular networks. Pathogen evolution has taken advantage of the cellular machineries mediating tSV budding. For instance, the HIV-1 Gag protein facilitates the transfer of viral information among interacting cells by hijacking components of the ESCRT system, mimicking the budding of SE [4, 56, 57]. We expect BSLBs will also help unravelling new pathogenic determinants hijacking vesicular messages across infected cell interfaces, broadening our understanding of ‘trojan horses’ driving infectious diseases [56].

## FIGURE LEGENDS

**Supplementary Figure 1; related to** **Fig. 1. Multiparametric analyses of released tSV show a highly heterogeneous population of vesicles being transferred at the IS.** (A) T cell: BSLB synapses and the release of synaptic ectosomes containing engaged T cell molecules can be studied by conventional multicolour flow cytometry. Shown is the gating strategy to discriminate single BSLB and T cells following conjugate dissociation, and the biparametric overlays of single BSLBs either null (orange), agonistic (brown) and non-stimulated (light green) and TCR-stimulated cells (dark green) for BST2, CD63, TCRαβ and CD38. (B) conventional FCM analyses of T cells after conjugate dissociation shows neligible uptake of fluorescent lipids by the stimulated T cells, which is observed across a broad range of α-CD3ε Fab densities. Data shown represent the % of the total Atto 565 signal observed in null BSLB. (C) Nanoparticle Tracking analysis size distribution of tSV and EVs isolated by differential centrifugation. (D) using the size calibration beads (top left panel), EVs (green and top middle panel) tSV (violet and top right panel) were compared across different bin sizes (SS-A range) as defined by the CV of standard bead population. (E) Size distribution measured by NanoFCM of TCR, CD63, CD40L, BST2 and CD81 positive EVs (green) and tSV (violet). (F) Example overlaid histograms showing the comparison of TCRαβ^+^ vesicle size using benchmark calibration silica beads (grey). Compared TCRαβ^+^ EVs (yellow) and tSV (blue) were isolated by differential and ultracentrifugation as described in Methods. (G) NanoFCM for the comparison of TCRαβ expression in EVs and eluted tSV. Expression was normalised to the signal measured in isotype-labelled controls (TCRαβ MFI / Isotype MFI). (H) Immunoblot showing the size comparison between tSV-associated CD40L and a recombinant human CD40L ectodomain. Null BSLB were loaded only with ICAM1 and CD40. Normality was determined using Shapiro-Wilk test and statistical significance was determined by Multiple paired t-test corrected by two-stage step-up method of Benjamini, Krieger and Yekutieli (FDR Q= 5%)(D), by unpaired double tailed t-test (E) and by unpaired, two-tailed t-test (G). *P* values are indicated above bars for the relevant comparisons. Data represent means ± SEM from n=4 independent experiments.

**Supplementary Figure 2; related to** **Fig. 1. TIRFM and Airyscan confocal imaging of T cell synapses indicate a high spatial segregation of T cell tSV.** (A) Pearson’s correlation coefficient (PCC) for the spatial colocalization between CD40L and other vesicle markers released within the cleft of TH synapses. Each dot represents the average PCC per synapse with representative TIRFM images shown in B-E. Glycans in the PM and tSV are labelled with Wheat Germ Agglutinin (WGA). SIRC: surface interference reflection contrast microscopy. Scale bar = 5 µm. (B-E) Representative TIRFM microphotographs showing the discrete spatial distribution of CD40L and other extracellular vesicle markers in non-permeabilised, surface-stained TH synapses stained with monoclonal antibodies specific to CD40L and CD63 (B); CD81 (C); BST2 (D); and ADAM10 (E, as control). F-I) Airyscan confocal microphotographs showing the 3D spatial distribution of CD40L and other tSV markers, namely (F) CD63, (G) BST2, (H) TCRαβ, and (I) CD81 on synapses formed between TH and glass-SLB after 15 min of interaction. As a counterstaining we used WGA CF594, which labels cell surface glycans. Right panels: orthogonal views of TH synapses with the Z-plane spanning the synaptic cleft. Scale bar = 5 µm. In A-I, TH were stimulated on glass-supported lipid bilayers containing 200 molec./µm^2^ of ICAM1, 20 molec./µm^2^ of CD40 and 30 molec./µm^2^ of α-CD3ε Fab. In (A) Normality was determined using Shapiro-Wilk test and statistical significance was determined by Kruskal-Wallis test corrected for multiple comparisons (Dunn’s). Data representative of n=2 independent experiments. *p≤0.05, **p≤0.005, ***p≤0.0005, ****p≤0.0001 and ns = not significant.

**Supplementary Figure 3; related to** **Fig. 1. The output from different subcellular compartments underpins tSV heterogeneity**. TH were preincubated for 30 min with a panel of 12 inhibitors and then co-cultured with control and agonist BSLB at a 1:1 ratio while keeping inhibitor concentrations. When possible, we used compounds redundantly targeting subcellular processes to elucidate effects related with the shared targets rather than to off-targets and used concentrations validated elsewhere [20, 21, 58–78] (Table S1). (A) After 90 min of culture, we measured the cell surface expression of key synaptic and tSV markers, and (B) the stability of cellular contacts by tracking the percent of remaining cell: BSLB conjugates following cold incubation. BSLB presenting 200 molec./µm^2^ ICAM1, 20 molec./µm^2^ of CD40 and increasing densities of anti-CD3ε Fab were used in all experiments. (C-F) Transfer of TCRαβ^+^ (C), CD40L^+^ (D), BST2^+^ (E), and CD63^+^ (F) tSV to BSLB compared and normalised to the maximum transfer observed in control untreated cells (i.e., maximum measured transfer (T_max_) per donor and at 2,000 molec./µm^2^). Right heatmaps display *P-*values for the comparison of tSV transfer between controls and inhibitors, and across different α-CD3ε-Fab densities. (G-J) Dynamics of TCRαβ (G), CD40L (H) BST2 (I) and CD63 (J) cell surface expression normalised to the baseline of untreated controls. In (J) out of range values (>200% are represented in orange). Statistical significance and *P*-values were determined using mixed-effects model ANOVA with Geisser-Greeenhouse correction for the determination of normalised protein expression across different donors (A), and multiple t-test to compare frequency of conjugates (B), levels of T_max_% tSV transfer (C-F), and levels of normalised cell surface expression of relevant markers as % of control baseline (100%) (G-J) across different α-CD3ε Fab densities. *p≤0.05, **p≤0.005, ***p≤0.0005, ****p≤0.0001 and ns = not significant. N=6 different donors.

**Supplementary Figure 4; related to** **Figure 2**. (A-C) Isolation and expansion of FoxP3^+^-enriched cCD4^+^ T cell cultures (termed herein as Treg). (A) sorting of cells on day 0 based on the high expression of CD4, negative expression of CD127 and further separation of CD25 low and the 5% highest population for CD25. Sorting purity controls are shown. Cells were then expanded for 14 days and used for experimentation between day 15 and day 17. (B) Example of intracellular staining for FoxP3 on day 15, and (C) the quantification of percent FOXP3^+^ cells on day 15 of culture. (D-I) BSLBs can also be tailored for the study of CART cell tSV. Data shown in A-E represent means +/- SEM from two independent experiments evaluating the transfer of the chimeric antigen receptor (CAR) (D), the TCR co-receptor CD8(E), CD63(F), Perforin(G), and as its comparison to TCRαβ at the tested densities (H). In D-H, Normality was determined using Shapiro-Wilk test and statistical significance was determined by Multiple t-test. *p≤ .05, **p≤0.005, ***p≤0.0005, ****p≤0.0001 and ns = not significant. (I) Representative pseudocolor plots showing the gating strategy for identifying single BSLBs and single CART. (D-G) Representative half-overlaid histograms are shown to the right. The differences in synaptic transfer at comparable α-CD3ε and HLA-A2: NY-ESO-1 densities (140 molec./µm^2^). tSV transfer instigated by either the CAR antigen (top panels; teal BSLBs), or the α-CD3ε Fab (bottom panels, violet BSLBs) are shown for each marker. Cells are shown in dark blue (no antigen/α-CD3), and light blue (co-cultured with either HLA-A2: NY-ESO-1 or α-CD3ε). Light grey BSLBs represent those loaded with no antigen. (J-K) Increased synaptic transfer of CD39 (J) and CD73 (K) to BSLB associates with higher central clustering of CD39 and CD73 in Treg synapses imaged by TIRFM. Left panels: TIRFM images comparing TH (top) and Treg (bottom) showing the distribution of the ectonucleotidases (A) CD39 and (B) CD73 in mature ISs. Middle panels show background subtracted mean fluorescence intensities (corrected MFI) for CD39 and CD73 in the contacts of TH and Treg cells stimulated on planar glass SLB containing 200 molec./µm^2^ of ICAM1, 20 molec./µm^2^ of CD40 and 30 molec./µm^2^ of α-CD3ε Fab (clone UCHT-1). SIRC: surface interference reflection contrast microscopy. Cell surface glycans are labelled with Wheat Germ Agglutinin (WGA). White arrowheads show released WGA^+^ CD73^+^ vesicles (diffraction limited). Scale bar = 5 µm. Two independent experiments are shown. Points represent individual synapses. Right panels: transfer behavior to BSLB relates to increased surface expression of ectonucleotidases in Treg cells (blue) compared to TH (green). Normality was determined using Shapiro-Wilk test and statistical significance was determined by two-tailed Mann-Whitney unpaired test. *p≤0.05, **p≤0.005, ***p≤0.0005, ****p≤0.0001 and ns = not significant.

**Supplementary Figure 5; related to** **Fig. 4**. (A) Surface protein expression for TH from 9 different donors edited using CRISPR/Cas9 targeting CD4, CD81, ADAM10, BST2, and TSG101. (B) Representative half-overlaid histograms and bi-parametric contour plots showing the surface expression levels of edited proteins (headings) and their relationship with the expression of CD40L on cells (Y-axis). (C) Relationship between CD40L expression and CD3ξ expression *CD4*-(teal) and *ADAM10*-edited cells (orange). (D) TSG101 downregulation resulting from CRISPR/Cas9-genome editing shown normalized to beta-actin (left panel). (E-J) Dynamics of tSV transfer between different CRISPR/Cas9 edited cells and the resulting downregulation of the markers on the cell surface. Non-linear, three-parameter regression analyses (F-test) comparing TCRαβ^+^ and CD40L^+^ tSV transfer among CRISPR-Cas9 knockdowns and in response to increasing densities of agonistic α-CD3ε-Fab. (E) T_max_ for TCR^+^ and CD40L^+^ tSV transfer to BSLB among different CRISPR/Cas9-edited cells. Data represent means normalised to *CD4*-edited controls. (F) α-CD3ε Fab EC_50_ for TCR^+^ and CD40L^+^ tSV transfer to BSLB among different CRISPR/Cas9-edited cells. Data represent means normalised to *CD4*-edited controls. (G) TCRαβ^+^ tSV (as TCRαβ^+^ molec./BSLB) transfer to BSLB at increasing α-CD3ε Fab densities, and (H) its resulting downregulation on the surface of T cells (H). (I) CD40L^+^ tSV (as CD40L^+^ molec./BSLB) transfer to BSLB at increasing α-CD3ε Fab densities, and (J) its resulting upregulation on the surface of T cells following stimulation with α-CD3ε-Fab.

**Supplementary Figure 6; related to** **Fig. 5**. (A) Sorting of BSLB after conjugate dissociation for the sequencing of tSVs small RNA content. Upper panel: Pseudocolor plots showing the gating strategy for sorting single BSLB for the downstream sequencing of small RNAs. Shown are post-sorting controls. Right overlaid histograms showing TCRαβ on BSLB loaded with α-CD3ε Fab (violet) versus null controls (grey) after sorting from T cell: BSLB co-cultures. Bottom panel: post-sort control showing the enrichment of single cells. (B) KEGG GO pathway enrichment analyses reveals unique miR to both T-cell tSV and EVs and sharing association with common biological processes. Heatmaps detailing miR species and KEGG GO pathways functionally enriched in TH tSV, TH EVs, CTL tSVs and CTL EVs; Heatmap values represent log10 p-values. (C-F) Bubble plots depicting enriched target pathways associated with miR identified in TH tSV (C), TH EVs (D), CTL tSVs (E), and CTL EVs (F). Circles represent different GO terms, and circle size represents the number of targets associated with the given GO category. The overall colour represents the cluster (a broader category) encompassing the enriched GO categories. (G-H) Number of miR targets across different GO pathways. CTL tSV and EVs (G), and TH tSVs and EVs (H) are compared. Shared miR targets per GO category are shown in grey, unique targets for tSVs are shown in magenta and unique targets for EVs miR are shown in green.

**Supplementary Figure 7; related to** **Fig. 5. SVs are enriched in RBPs, which localise as discrete puncta in the synaptic cleft of mature TH ISs.** (A) LC-MS/MS analyses of enriched proteins found in eluted tSVs from a-CD3 coated BSLB and null BSLB containing only ICAM1 and CD40. (B-C) differentially expressed proteins in tSV were then used for the differential expression analyses of tSV with EVs from TH (B) and CTL (C). (D-G) Enriched proteins in tSV and EVs were identified by averaged-based normalisation of peptide intensities (log10). (D) tSV and EVs showing the differential enrichment of contaminating proteins. (E) . (F) GSEA using PANTHER [79] revealed a higher enrichment of nucleic-acid binding proteins in the LC-MS/MS of tSV (magenta) derived from TH (left) and CTL (middle left) as compared with syngeneic EVs (green) of TH (middle right) and CTLs (right). Values represent the fold enrichment of identified proteins per GO term. (H) Venn-Diagram showing the co-identification of RBPs in LC-MS/MS datasets generated by two different isolation methods, namely tSV elution and sorting of BSLB. (I) TIRFM imaging of the RBP YBX1 in the SLB-contacting pole of T cell IS after 20 minutes of interaction with SLBs. YBX1 was enriched in TH tSV as defined by analyses in D and in new and previously published[5] proteomics from isolated ectosomes. T cells were fixed, permeabilised, and stained with monoclonal antibodies against vimentin for the identification of the pSMAC (yellow) and the cSMAC (dark central area). White arrows indicate the localisation of YBX1 in vesicular tracks of cells resuming migration and in the cSMAC -synaptic cleft. (J) as in I, T cells were stained against SF3B3[5]. (K) A rabbit monoclonal IgG isotype control was used as a control. (L-M) TH cells were pre-stained with BODIPY™ TR Ceramide and SYTO™ RNASelect™ to track the polarisation and localisation of ceramide and RNA enriched structures in the synaptic cleft. (L) Examples of some T cell kinapses showing the release of ceramide^+^ and RNA^+^ vesicles in the tracks of cells resuming migration (yellow arrows). (M) *Left:* Example of a TH synapse showing localisation of ceramide^+^ and RNA^+^ puncta in transit to the cSMAC. *Right*: We used the super-resolution provided by eTIRF-SIM to visualise the arrangement of RNA^+^ puncta at 5 min of synapse assembly. Most puncta associate to ceramide enriched ER-like structures lining the cSMAC (synaptic cleft, yellow arrows), suggesting the rapid polarisation of RNA transport machineries to the boundary of the synaptic cleft. Scale bars = 5 µm. Imaging representative of at least three independent experiments and four different donors.

## ACKNOWLEDGEMENTS

We are grateful to our laboratory members and the Kennedy Institute of Rheumatology community for constructive scientific discussions, especially to James Felce, Jonathan Webber, Štefan Balint, Alexander Mørch and Kristina Correa. We thank the technical support of Heather Rada, Kellie Johnson and Ekaterina Zvezdova (the latter two from BioLegend). We thank Professor Catarina E. Hioe for kindly providing the HIV-1 gp120 protein. We would also like to thank all the anonymised blood donors who contributed to our study. This work was funded by Wellcome Trust Principal Research Fellowship 100262Z/12/Z, the ERC Advanced Grant (SYNECT AdG 670930), and the Kennedy Trust for Rheumatology Research (KTRR) (all three to M.L.D.). P.F.C.D was supported by EMBO Long-Term Fellowship (ALTF 1420-2015, in conjunction with the European Commission (LTFCOFUND2013, GA-2013-609409) and Marie Sklodowska-Curie Actions) and Oxford-Bristol Myers Squibb Fellowship. A.K. was supported by H2020 and the Research Council of Norway (in conjunction with Marie Sklodowska-Curie Actions 275466; to A.K.). M.F. and H.C.Y. thank the Wellcome Trust (212343/Z/18/Z) and EPSRC (EP/S004459/1). The eTIRF-SIM platform was built in collaboration with Micron (www.micronoxford.com), an Oxford-wide advanced microscopy technology consortium supported by Wellcome Strategic Awards (091911 and 107457), and with additional funds from an MRC/EPSRC/BBSRC next-generation imaging award and the Kennedy Trust for Rheumatology Research through the Kennedy Institute Cell Dynamics Platform. We acknowledge the generous support of the Kennedy Trust for Rheumatology Research, IDRM and Carl Zeiss GMBH for the Airyscan LSM 980 confocal microscope used in this research. Y.P., T.D. and R.F. were supported by the Chinese Academy of Medical Sciences (CAMS) Innovation Fund for Medical Sciences (CIFMS), China (grant number: 2018-I2M-2-002) and UK Medical Research Council (MRC); E.S. was supported by Newton-Katip Celebi Institutional Links grant (352333122) and SciLifeLab fellowship (to E.S.). F.S-M. was supported by grants SAF2017-82886-R from the Spanish Ministry of Economy and Competitiveness (MINECO), and “La Caixa” Banking Foundation (HR17-00016). We thank the NIH Tetramer Core Facility for the synthesis of the HLA-DRB1*09:01 monomers used in this study. We thank the Oxford Genomics Centre at the Wellcome Centre for Human Genetics (funded by Wellcome Trust grant reference 203141/Z/16/Z) for the generation and initial processing of the sequencing data. Finally, we thank the MS laboratory at the Target Discovery Institute NDM (Oxford) led by Benedikt M. Kessler. Pablo F. Céspedes is also known as Pablo F. Céspedes-Donoso (https://orcid.org/0000-0002-1641-4107).

## MATERIAL AND METHODS

### Ethics

Different T cell populations were isolated from leukapheresis reduction system (LRS) chambers from de-identified, non-clinical healthy donors. The non-clinical issue division of the National Health Service and the Inter-Divisional Research Ethics Committee of the University of Oxford approved the use of LRS chambers (REC 11/H0711/7 and R51997/RE001).

### Isolation and expansion of human CD4^+^, CD8^+^ and Treg cells from peripheral blood

Briefly, human T cells were isolated from leukoreduction system (LRS)-concentrated peripheral blood by negative immunodensity selection using either CD4^+^, CD8^+^ or CD4^+^CD127^low^ (RosetteSep™, StemCell™ Technologies, #15022, #15023, and #15361). Enriched CD4^+^ CD127^low^ cells were immediately used for fluorescence-activated cell sorting of regulatory T cells (Treg) based on low CD127 fluorescence and high CD25 fluorescence. Briefly, Tregs were sorted from enriched CD4^+^ CD127^neg^ peripheral blood cells using a FACSAria III cell sorter. A nested gating strategy was used in which CD4^+^ CD127^neg^ CD25^high^ cells were gated such that only the brightest 5% of CD25^high^ cells were sorted. Recovered cells were stimulated with Human T-activator CD3/CD28 Dynabeads (ThermoFisher) at a bead to cell ratio of 3:1 and 1,000 U/mL of recombinant human IL-2 (Peprotech). Fresh IL-2 containing media was added every 48 h and Human T-activator CD3/CD28 Dynabeads were added again at day 7 and removed at day 15 of expansion. This protocol allowed us the enhanced recovery of FoxP3^+^ T cells by day 15 of expansion (see Fig. S4A-C). On the other hand, conventional CD4^+^ and CD8^+^ T cells were expanded using a bead to cell ratio of 1:1 and 100 U/mL of IL-2. After 3 days of activation, Human T-activator CD3/CD28 Dynabeads were removed and the conventional CD4^+^ and CD8^+^ cells kept in complete RPMI media supplemented with 100 U/mL of IL-2. All BSLB synaptic transfer experiments were performed after preconditioning of T cells to IL-2-depleted complete RPMI 1640 media containing 10% of either heat-inactivated AB human serum (Treg & controls) or foetal bovine serum, and 100 µM non-essential amino acids, 10 mM HEPES, 2 mM L-glutamine, 1 mM sodium pyruvate, 100 U/ml of penicillin and 100 µg/mL of streptomycin).

### Culture of CD4^+^ T cell clones

The HLA-DRB1*09:01-restricted T cell clones 16, 35 and 40 (all specific against the influenza H3 HA_338-355_ peptide: NVPEKQTRGIFGAIAGFI) were expanded using at a ratio of 1 clone: 2 feeder cells (irradiated, pooled PBMCs from 2-3 healthy donors) at a total cell concentration of 3 × 10^6^ cells/mL in RPMI 1640 supplemented with 10% heat-inactivated AB human serum and 30 µg/ml of PHA for three days. Then, 100 U/ml of recombinant human IL-2 were added to fresh media, which was replaced every 2 days. Clones were used between days 8 and 12 of culture.

### Production of chimeric antigen receptor (CAR) recombinant CD8^+^ T cells

The CAR constructs that bind the NY-ESO-1_157-165_ HLA-A2 complex were a kind gift from Cristoph Renner (Zurich, Switzerland) and described elsewhere [80, 81]. Briefly, the T1 CAR binds the C9V NY-ESO-1 APL with a K_D_ of ∼2 nM. The CAR construct was packaged within third-generation self-inactivating lentiviral transfer vectors with the EF1α promoter and the Woodchuck Hepatitis Virus Post-Transcriptional Regulatory Element. The CAR features a scFv binding domain, Ig domain spacers derived from human IgG1, the CD28 transmembrane domain and the CD3ζ signalling tail. For lentiviral production, 293T cells (ATCC CRL-3216) were co-transfected with a mix of the VSV-G (370 ng), the lentiviral CAR (800 ng), and RSV-Rev and GAG (950 ng each) plasmids. The plasmid mix was prepared in DMEM (Merck, #D6429) containing X-treme Gene HP DNA Transfection Reagent (Merck, #6366546001) and after 20 min preincubation at RT, the mix was added to 293T cells in a dropwise manner. Twenty-four h after isolation and stimulation with Human T-activator CD3/CD28 Dynabeads, CD8^+^ T cells were transduced with freshly harvested and 0.45 µm-filtered supernatant containing 50 U/mL of recombinant human IL-2. Three days after transduction Dynabeads were removed and the cells kept in culture at a concentration of 1×10^6^ cells/mL in IL-2 supplemented RPMI media (see above). Transduction efficiency was determined between days 8-10 of culture using a polyclonal goat anti-human IgG Fc PE (ThermoFisher Scientific, #12-4998-82) and the synaptic transfer to BSLB addressed immediately.

### Bead Supported Lipid Bilayers (BSLB)

Non-functionalized silica beads (5.00 ± 0.05 µm diameter, Bangs Laboratories, Inc.) were washed extensively with PBS in 1.5 ml conical microcentrifuge tubes. BSLBs were formed by incubation with mixtures of liposomes to generate a final lipid composition of 0.2 mol% Atto-DOPE; 12.5 mol% DOGS-NTA in DOPC at a total lipid concentration of 0.4 mM. In this work, we used DOPE lipids conjugated with either Atto 390, 488, 565 or 647. The resultant BSLB were washed with 1% human serum albumin (HSA)-supplemented HEPES-buffered saline (HBS), referred herein as HBS/HSA buffer. To saturate NTA sites, BLSB were then blocked with 5% casein 100 μM NiSO_4_ for 20 minutes. After two washes, BSLB were loaded with concentrations of His-tagged proteins required to achieve the indicated molecular densities (in range of 1-100 nM; please refer to each Figure legend). Excess proteins were removed by washing with HBS/HSA after 30 minutes. T cells (2.5 × 10^5^/well) were incubated with BSLB at 1:1 ratio in either U-bottomed or V-bottomed 96 well plate (Corning) for 90 min at 37°C in 100 µl HBS/HSA. For gentle dissociation of BSLB: cell conjugates, culture plates were gradually cooled down by incubation at RT for 15 min, followed by incubation on ice. After 45 min, cells and BSLB were centrifuged at 300 *g* for 5 min prior to resuspension in ice-cold 5% BSA in PBS pH 7.4. Single BSLB and cells were gently resuspended prior to staining for flow cytometry analysis or sorting.

### Multicolour Flow Cytometry (FCM)

Staining with fluorescent dye conjugated antibodies was performed immediately after dissociation of cells and BSLB conjugates. Staining was performed in ice-cold 5% BSA in PBS pH 7.4 (0.22 µm-filtered) for a minimum of 30 min at ^+^4°C. Then, cells and BSLB were washed three times and acquired immediately using an LSRFortessa X-20 flow cytometer equipped with a High Throughput Sampler (HTS). For absolute quantification, we used Quantum Molecules of Equivalent Soluble Fluorescent dye (MESF) beads (see below), which were first acquired to set photomultiplier voltages to position all the calibration peaks within an optimal arbitrary fluorescence units’ dynamic range (between 10^1^ and 2 × 10^5^, and before compensation). Fluorescence spectral overlap compensation was then performed using single colour-labelled cells and BSLB, and unlabelled BSLB and cells. For markers displaying low surface expression levels unstained and single colour stained UltraComp eBeads (Thermo Fisher Scientific Inc.; #01-2222-42) were used for the calculation of compensation matrixes. After application of the resulting compensation matrix, experimental specimens and Quantum MESF beads were acquired using the same instrument settings. In most experiments acquisition was set up such as a minimum of 5 × 10^4^ single BSLBs were recorded. To reduce the time of acquisition of high throughput experiments a minimum of 1 × 10^4^ single BSLBs were acquired per condition instead. All dye-conjugated antibodies described in this study are listed in Table S5.

### Calibration of flow cytometry (FCM) data

T cells and BSLB were analysed using antibodies with known fluorescent dye to Ab ratio (F/P) in parallel with the Quantum MESF beads (Bangs Laboratories, Inc. IN, USA), allowing the calculation of the absolute number of antibodies bound per T cell and per BSLB after subtraction of unspecific signals given by isotype control antibodies. We used MESF standard beads labelled with the Alexa Fluor® dyes 488 and 647 to estimate number of dye molecules from mean fluorescence intensities (corrected and geometric, cGMFI). Briefly, MESF beads provided 5 different populations of beads with increasing GMFI allowing the linear regression of corrected GMFI (cGMFI) over MESF. The resulting slope is then used for the interpolation of total fluorochromes bound to either BSLB or cells from cGFMI values. Since we also used antibodies with known fluorochromes per protein (F/P), we then estimated the number of bound antibodies (and hence molecules) per BSLB by dividing the estimated fluorochrome number by the detection antibody F/P value (Number of molec./event = Fluorescent molec._event_/(F/P)_Ab_).

### Trans-synaptic vesicle elution from BSLBs

Cells and BSLBs were incubated at a 1:1 ratio for 90 min at 37 °C and 5% CO_2_ in Phenol Red-free FBS-free RPMI 1640 supplemented with 100 µM non-essential amino acids, 2 mM L-glutamine, 1 mM sodium pyruvate, 100 U/ml of penicillin and 100 µg/mL of streptomycin. Cultures needed to be scaled up and therefore a CO_2_ incubator and supplemented FBS-free, Phenol Red-free RPMI 1640 was used instead of 1 %HSA/HBS. Cells and BSLBs were collected at different time points of the elution protocol to control with FCM both the transfer of material from T cells to BSLB, and the release of tSV from BSLB upon addition of EDTA for the chelation of Ni. Briefly, elution was performed as follows; first conjugates were cooled down 15 min at RT and then 40 min on ice to gentle separate cells and BSLB. Then, conjugate suspensions were resuspended by adding 3 volumes of ice-cold PBS pH 7.4 and centrifuged at 100*g* for 1 min and ^+^4°C to enrich for BSLB. After two washes with ice cold PBS pH 7.4, BSLB were resuspended in ice-cold 50 mM EDTA in PBS pH 7.4 for 2h. BSLB-free supernatants were then centrifuged at ^+^4°C in sequential steps at 300*g* for 5 min (twice), then at 2,000*g* for 10 min and 10,000*g* for 10 min. Finally, recovered supernatants were centrifuged at 120,000*g* for 4h at ^+^4°C and the pellets washed once more with PBS pH 7.4 and kept at ^+^4°C or frozen at -80 °C until analyses by either NTA, NanoFCM or immunoblotting. Steadily released EVs were isolated by the same differential centrifugation procedure indicated above. To minimize serum and debris contamination, T cells were cultured in OptiMEM-I supplemented with 100 U/mL of IL-2, 100 µM non-essential amino acids, 2 mM L-glutamine, 1 mM sodium pyruvate, 100 U/ml of penicillin and 100 µg/mL of streptomycin for no longer than 48 h. All samples isolated by UC were resuspended in 0.22 µm-filtered PBS pH 7.4 to reduce background signals in downstream particle measurement analyses with either nanoFCM or Nanoparticle Tracking Analysis (NTA).

### TEM

The negative staining of thinly spread vesicle preparations for transmission electron microscopy (TEM) was performed as described elsewhere [82]. Briefly, carbon support film-coated 3 mm copper grids (300 mesh) were plasma treated for 20 s using a Leica EM ACE200 Vacuum Coater. Then, 10 µL of isolated vesicle populations were deposited on and incubated at RT for 5 min, followed by removal of excess sample with a Whatman N°1 paper and staining with 2% uranyl acetate for 10 s at RT. After removal of excess uranyl acetate the samples were dried for 10 min and analysed using a Tecnai 12 TEM at 120 kV using a Gatan OneView CMOS camera. A final magnification of 29,000x was used for imaging of isolated vesicle populations.

### NTA

Eluted tSV and purified EV preparations were resuspended in 0.22 µm-filtered PBS pH 7.4 in a 1:100 dilution and kept on ice for Nanoparticle Tracking Analysis (NTA). The instrument used for NTA was Nanosight NS300 (Malvern Instruments Ltd) set on light scattering mode and instrument sensitivity of 15. Measurements were taken with the aid of a syringe pump to improve reproducibility. Three sequential recordings of 60 seconds each were obtained per sample and NTA 3.2 software was used to process and average the three recordings to determine the mean size.

### Nano Flow Cytometry

Flow NanoAnalyzer model type N30E (NanoFCM Inc., Xiamen, China) that allows single exosomes detection was used to determine the size distribution and granular concentration of E Vs. The Nano-flow cytometry analysis was performed using the Flow NanoAnalyzer (NanoFCM Co., LTD) according to manufacturer’s instructions. The Silica Nanospheres Cocktail (S16M-Exo, NanoFCM) was employed as the size standard to construct a calibration curve to allow the conversion of side scatter intensities to particle size. A concentration standard (200nm PS QC beads, NanoFCM) was used to measure particle concentration. The laser used was a 488nm laser at 25/40 mW, 10% ss decay. Lens filters equipped were 525/40 (AF488) and 580/40 (PE). Before staining samples were acquired to determine particle concentration such that a total of 10^8^ vesicles of either tSV or EVs were labelled per condition. Before use, fluorochrome-conjugated antibodies were spun down at 10,000g for 10 min at +4°C. Isotype control antibodies were used at the same effective concentrations (ranging from 0.2 to 5 µg/mL) and incubation times. Staining antibodies were conjugated to AF488 and AF647. Stainings were performed for 30 min on ice. Samples were washed and centrifuged for 1 h at 100,000*g* and +4°C. Labelled vesicles were then resuspended in 50 µL of PBS. Buffer alone (PBS), unstained vesicle, isotype controls and autothresholding were used to define positivity.

### Planar Supported Lipid Bilayers (PSLB)

Liposome mixtures were injected into flow chambers formed by sealing acid piranha and plasma-cleaned glass coverslips to adhesive backed plastic manifolds with 6 flow channels (StickySlide VI 0.4; Ibidi). After 30 minutes the channels were flushed with HBS-HSA without introducing air bubbles to remove excess Liposomes. After blocking for 20 min with 5% casein supplemented with 100 µM NiCl_2_, to saturate NTA sites, followed by washing and then His-tagged proteins were incubated on bilayers for additional 30 min. Protein concentrations required to achieve desired densities on bilayers were calculated from calibration curves constructed from FCM measurements on BSLB and analysed alongside MESF beads (MESF; Quantum™ Bangs Laboratories Inc.). Bilayers were continuous liquid disordered phase as determined by fluorescence recovery after photobleaching with a 10 µm bleach spot on an FV1200 confocal microscope (Olympus).

### Immunological Synapse formation on glass-SLB

Prior to immunological synapse imaging, primary T cell blasts were washed twice and resuspended in prewarmed HBS/HSA buffer to a final concentration of 5 × 10^6^ cells/mL. Then, 5 × 10^5^ T cells were stimulated for 20 min at 37°C and 5% CO_2_ over PSLB containing 30 molec./µm^2^ anti-CD3ε Fab (clone UCHT-1; Alexa Fluor® (AF) 488 or unlabelled), 200 molec./µm^2^ of ICAM1 AF405, and 100 molec./µm^2^ of CD40 (AF488 or unlabelled). Then, cells were stained with 1 µg/mL of wheat germ agglutinin (WGA) conjugated to CF568 (Biotium; #29077-1) and 1 µg/mL anti-CD154 (CD40L) clone 24-31 AF647 for 15 min at RT and in the dark. Cells were washed three times with 5% BSA 20 mM HEPES 2 mM MgCl_2_ in PBS and fixed with pre-warmed 4% electron microscopy grade formaldehyde in PBS pH 7.4 containing 2 mM MgCl_2_ for 10 min at RT and in the dark. After two washes, cells were kept in HBS/HSA until imaging by TIRFM. For intracellular staining, cells were permeabilized for 3 min with 0.1% Triton X-100 in PBS, washed and blocked for 60 min with 5% BSA in PBS before staining with primary antibodies (1 µg/mL for 1h at RT). Primary antibodies included Rabbit anti-human TSG101 clone EPR7130B AF647 (#ab207664), anti-human YB1 (YBX1) clone EPR22682-2 (#ab255606), anti-human SF3B3 clone EPR18440 (#ab209402), and Rabbit Isotype control EPR25A AF647 (#ab199093). After three washes, cells were blocked with 0.22 µm-filtered 5% Donkey Serum for 1 h at RT before staining for 1h with AF647 AffiniPure F(ab’)□ Fragment Donkey Anti-Rabbit IgG (H+L) (Jackson ImmunoResearch, #711-606-152). After four washes, cells were washed and then stained with anti-Vimentin clone EPR3776 AF555 (#ab203428). Cells were washed four times before acquisition.

### TIRFM

TIRFM was performed on an Olympus IX83 inverted microscope equipped with a 4-line (405 nm, 488 nm, 561 nm, and 640 nm laser) illumination system. The system was fitted with an Olympus UApON 150x 1.45 numerical aperture objective, and a Photometrics Evolve delta EMCCD camera to provide Nyquist sampling. Quantification of fluorescence intensity was performed with Fiji/ImageJ (National Institute of Health) and MATLAB R2019b. A batch measure macro was used to automatically segment cell:SLB contacts based on internal reflection microscopy followed by both background subtraction and the measure of fluorescence across different channels.

### Airyscan microscopy

Airyscan imaging of T cell and SLB was performed on a Zeiss Axio Observer.Z1 LSM 980 confocal laser-scanning microscope equipped with an Airyscan 2 module (Zeiss, Oberkochen, Germany) consisting of 32 concentrically arranged GaAsP PMT detectors and 2 MA-PMT channels. The acquisition was performed using the Airyscan super-resolution (SR) and best signal Smart Setup and a C Plan-Apochromat 63x/1.4 NA Oil DIC magnification objective. Illumination was provided by a Solid-State Light Source Colibri 7 LED lamp and by Diode lasers at 639 nm, 594 nm, and 488 nm with 0.4% laser power and 850V detector gain for all channels. The imaging field was defined using a 6.0X scan zoom (crop area) and a Z-coverage spanning the totality of the synaptic cleft as defined by WGA staining. The final acquisition settings included a sequential acquisition in the order 647/594/488, a frame size of 528 × 528 px, a pixel time of 7.95 µs, a pixel size of 0.043 × 0.043 × 0.16 µm, and a doubled pixel sampling with bidirectional mean intensity averaging of acquisition lines. Analyses were performed using the ZEN 3.2 system blue edition (Carl Zeiss Microscopy GmbH), and Fiji v2.1.0/1.53c (build 5f23140693)[83].

### eTIRF-SIM

A custom-built eTIRF-SIM microscope system was used and detailed elsewhere[84]. Structured illumination was obtained via a grating pattern generated by a ferroelectric spatial light modulator (SLM, Forth Dimension Displays, QXGA3DM). After, the first diffraction orders are selected by the mask and sent to the Olympus IX83 microscope head. The distance between diffraction orders were tuned by SLM settings ultimately defining the illumination angle and therefore the TIRF depth. Excitation wavelengths of 488 nm, 560 nm, and 640 nm were used (MPB communications Inc., 500mW, 2RU-VFL-P-500-488-B1R, 2RU-VFL-P-500-560-B1R). Sample illumination was carried out using a high-NA TIRF objective (Olympus Plan-Apochromat 100X 1.49NA). The emitted fluorescence was collected by the same objective and sent onto sCMOS cameras recording the raw data (Hamamatsu, Orca Flash 4.0 v2 sCMOS). The excitation numerical aperture (NA) was adjusted for each wavelength by changing the period of the grating pattern at the SLM, which allows controlling the TIRF angle and, therefore, the penetration depth of the evanescent wave. To achieve TIRF-SIM illumination at the interface between the cells and SLB, both excitation lights were sent with an incident NA ranging from 1.38 to 1.41. Prior to experiment acquisition alignment was perform by imaging 100 nm Tetraspeck fluorescent beads in all three excitation colours. Then, using Fiji plugin MultiStackReg – the beads images were used to adjust images and compensate for chromatic aberrations. A total of 9 raw images were acquired per frame and for a single excitation wavelength before switching to the next wavelength. Then, raw images were processed and reconstructed into SIM images by custom made software or ImageJ fairSIM plugin. All experiments were performed at physiological conditions using a micro-incubator (H301, Okolabs, Italy) at 37 °C and 5% CO_2_. For each frame, we used an acquisition time between 20 and 300 ms depending on the fluorescence signal levels and 2 colours, 18 frames total) every 0.4–5 s. One colour super-resolved image was reconstructed from 9 raw image frames (3 angles and 3 phases) using a reconstruction method described previously [85, 86].

### CRISPR/Cas9 genome editing

After 48 h of stimulation with Human T-activator CD3/CD28 Dynabeads (ThermoFisher) CD4^+^ lymphoblasts were recovered for transfection of CRISPR/Cas9 nucleoprotein complexes. Briefly, tracer RNA and crRNAs were mixed at equimolar concentrations (200 µM) and incubated at 95°C for 15 min. After cooling the gRNA mix down to RT, the Cas9 enzyme was added at final 20 µM (IDT, #1081061). After 15 min incubation at 37°C, electroporation enhancer was added to CRISPR-Cas9 nucleoprotein complexes following manufacturer’s guidelines, and immediately mixed with 1.5 × 10^6^ cells. Cells were then transferred to a 2-mm cuvette (Bio-Rad) and electroporated at 300V for 2ms using an ECM830 Square Wave electroporator (BTX). Immediately after transfection, cells were recovered with pre-warmed, IL-2 supplemented RPMI 1640 media and expanded for another 6 days. Synaptic transfer experiments to BSLB or planar SLB were performed on day 7 or 8 of culture and protein expression controls were carried out in parallel. All crRNAs sequences are summarised in Table S6. All on- and off-target scores were optimised by the crRNA supplier (IDT).

### Western Blotting

Whole cell lysates (WCL) were prepared by resuspending cell pellets in RIPA lysis and extraction buffer (Thermo Fisher Scientific, #89901) containing a Protease/Phosphatase inhibitor cocktail (Cell Signaling Technology (CST); #5872) to a final concentration of 2 × 10^7^ cells/mL. After sonication at +4°C (10 cycles of 30 s on/30 s off), WCL were centrifuged at 10,000*g* for 10 min at ^+^4°C, and the supernatants collected, mixed with loading solution and denatured at ^+^95°C for 10 min. For immunoblotting of BSLB eluted material, after centrifugation at 120,000*g* for 4 h at ^+^4°C, pellets were resuspended in RIPA lysis buffer containing a Protease/Phosphatase inhibitor cocktail, mixed with loading solution and denatured at ^+^95°C for 10 min. When indicated, lysed eluates and EVs from an equivalent number of originating cells were resolved to compare different vesicle populations. As positive control we used WCL equivalent to 2.25 × 10^5^ CD4^+^ lymphoblasts. Similarly, for CRISPR/Cas9 edited cells, WCL equivalent to 3 × 10^5^ cells were used per lane. Samples were resolved using 4-15% Mini-PROTEAN SDS-PAGE gel (Bio-Rad; #4561084), transferred to 0.45 µm nitrocellulose membranes (Bio-Rad, #1620115), blocked and incubated with the following primary antibodies: rabbit anti-human CD40 Ligand clone D5J9Y (CST, #15094), rabbit anti-GAPDH clone D16H11 (CST, #5174), mouse anti-β-actin clone 8H10D10 (CST, #3700), and rabbit anti-TSG101 clone EPR7130B (#ab125011). Then, membranes were incubated with IRDye® 680RD donkey anti-mouse IgG (H+L; LI-COR, #926-68072) and IRDye® 800CW donkey anti-rabbit IgG (H+L; LI-COR, #925-32213) secondary antibodies following manufacturer guidelines. After four washes, membranes were imaged and analysed using the Odyssey® CLx Near-Infrared detection system equipped with the Image Studio™ Lite quantification software (LI-COR, Lincoln, NE).

### RNA sequencing and miR analyses

Total cell RNA was extracted using the miRNeasy Tissue/Cells Advanced Mini Kit (Qiagen, #217604). Purified RNA yields and quality were assessed via Agilent 2100 Bioanalyzer using the Agilent 6000 RNA Pico chips (Agilent Technologies, # 5067-1513). Before library preparation the material was further quantified using RiboGreen (Invitrogen) on the FLUOstar OPTIMA plate reader (BMG Labtech) and the size profile and integrity analysed on the 2200 or 4200 TapeStation (Agilent, RNA ScreenTape). Input material was normalised to 200 ng or maximum mass for input volume prior to library preparation. Small RNA library preparation was completed using NEBNext Small RNA kit (NEB) following manufacturer’s instructions and applying the low input protocol modifications. Libraries were amplified (15 cycles) on a Tetrad (Bio-Rad) using in-house unique dual indexing primers as described elsewhere [87]. Size selection was performed using Pippin Prep instrument (Sage Science) using the 3% Agarose, dye free gel with internal standards (size selection: 120 to 230bp). Individual libraries were normalised using Qubit, and the size profile was analysed on the 2200 or 4200 TapeStation. 10 nM libraries were denatured and further diluted prior to loading on the sequencer. Two runs of single end sequencing were performed; run one was performed in an Illumina HiSeq 2500 system (1×60) and run two was performed using Illumina NextSeq 500/550 v2.5 Kits (75 cycles). Quality control and processing of the raw sequencing data was performed using sRNAbench [88] and miRQC [89] which allowed the assessment of sequencing yield, quality, percentage of miR-mapping reads, read length distribution and relative abundance of fragments from other RNA species. Functional and biological pathway enrichment analyses were performed on annotated miR species shared between the two independent sequencing runs and enriched in each EV category (i.e., SV and sEV). MIENTURNET and the KEGG annotation database were used. Statistical analysis for functional and biological pathway enrichment analyses using MIENTURNET were carried out by calculating the *P*-value resulting from the hypergeometric test. Motif analyses of bulk, enriched miR sequences in each sample were performed using MEME (Multiple Em for Motif Elucidation 5.1.0) [42]. To dissect the miR-target gene interactome of our enriched miR we used miRNet 2.0 [90] by first mapping input miR to the miR interaction knowledgebase comprising annotations from miRbase, miTarBase and ExoCarta together with interactions with other miR, genes and transcription factors from TransmiR 2.0, ENCODE, JASPAR and ChEA. The output of the ’miRs’ module of miRNet 2.0 provided the motif miR-target gene interactome enrichment analysis. We used FunSet [91] to graphically plot gene ontology (GO) enrichment analyses in 2D plots clustering GO terms based on semantic similarities and extracting representative terms for each cluster. The result of GO enrichment analysis using Funset was filtered by setting the FDR threshold to 0.05 (using the Benjamini-Hochberg procedure) and the *P*-values were calculated using the hypergeometric test.

### Mass spectrometry

Samples isolated by differential centrifugation were concentrated using 100,000 NMWL centrifugal filters (Amicon® Ultra, #UFC510024, Merck Millipore Ltd.) and prepared in S-Trap™ spin columns (#C02-micro-80) following manufacturer recommendations. Briefly, samples were reduced with 5 µL of 10 mM TCEP and alkylated with 50 mM of IAA for 30 min each, then acidified with 12% phosphoric acid 10:1 vol:vol, and then transferred to S-trap columns. Then, samples were precipitated using 1:8 vol:vol dilution of each sample in 90% methanol in 100 mM TEAB. Samples were then washed three times with 90% methanol in 100 mM TEAB. Samples were then resuspended in 50 µL of 50 mM TEAB and digested with trypsin (Promega, #V1115) overnight at 37°C. Peptides were eluted from the S-Trap by spinning for 1 min at 1,500*g* with 80 µL of 50 mM ammonium bicarbonate, 80 µL 0.1% FA and finally 80 mL of 50% ACN 0.1% FA. The eluates were dried down in a vacuum centrifuge and resuspended in 2% ACN 0.1% TFA prior to off-line high-pH reversed-phase fractionation using RP-S cartridges pre-primed with 100 µL ACN at 300 µL/min and equilibrated with 50 µL of 2% ACN 0.1% TFA at 10 µL/min. Samples were loaded at 5 mL/min and divided into 8 fractions (elution steps), which were run individually. Elutions were performed with increasing concentrations of 90% ACN, pH 10 in water, including final 5%, 10%, 12.5%, 15%, 20%, 22.5%, 25% and 50%. Fractions were further dried down in a vacuum centrifuge and resuspended in loading buffer. For LC-MS/MS acquisition 50-80 ng peptides were loaded onto preconditioned Evotips containing 0.1% FA in water. Preconditioning was performed by pre-priming of isopropanol-soaked tips with 20 µL of ACN 0.1% FA, following by centrifugation for 1 min at 700*g*, equilibration in water 0.1% FA and a final centrifugation for 1 min at 700*g*). Samples were run on a LC-MS system comprised of an Evosep One and Bruker timsTOF Pro. Peptides were separated on an 8 cm analytical C18 column (Evosep, 3 µm beads, 100 µm ID) using the pre-set 60 samples per day gradient on the Evosep One. Acquisition was done in PASEF mode (4 PASEF frames, 3 cycles overlap, oTOF control v6.0.0.12) including an ion mobility window between 1/k0 start=0.85 Vs/cm2 to 1/k0 end = 1.3 Vs/cm^2^, a ramp time of 100 ms with locked duty cycle and a mass range of 100-1,700 m/z. For proteomic analyses the raw files were searched against the reviewed Uniprot *Homo sapiens* database (retrieved 2,01,80,131) using MaxQuant version 1.6.10.43 [92] and its built-in contaminant database using tryptic specificity and allowing two missed cleavages.

### Statistical Analyses

Normality tests were performed using either Shapiro-Wilk or Kolmogorov-Smirnov tests. Statistical significance was determined by multiple comparisons performed either by one-way analysis of variance (ANOVA) or Kruskal-Wallis (corrected by Holm-Sidak or Dunn’s, respectively). Non-linear regressions using three or four parameters F-tests were also performed when indicated. All statistical tests were performed in GraphPad Prism v8 and are detailed in each figure legend. Means of significance are as follows: *p < 0.05, ** p < 0.005, ***p < 0.0005, and ****p < 0.0001.

## Notes

### Competing Interest Statement

Ben Peacock, Alice Law, and Dimitri Aubert are employed by NanoFCM Co., Ltd.

### Summary of Updates

We have amended some minor in-text errors and corrected Figure S7 annotations. We changed the terms synaptic vesicles to trans-synaptic vesicles (tSV) to avoid nomenclature conflicts with neuroscience. We also changed "steadily shed EVs" to "steadily released EVs" to follow ISEV standards. By doing so we are also referring to EVs as a broad population and not exclusively to those shed from the plasma membrane. We improved the analysis and representations of our LC-MS/MS data. More specifically, we used the material from null BSLBs (as noise) to clean our MS/MS datasets of tSV before comparing them with syngeneic EVs. The conclusions remain unchanged. We also updated the author list to acknowledge the contributions of all scientists making this work possible. We greatly appreciate all the comments from peers, which have improved the quality of our manuscript.

