## Supplementary figures and images for "Synthetic antigen-presenting cells reveal the diversity and functional specialisation of extracellular vesicles composing the fourth signal of T cell immunological synapses"

### Supplemental Figure 1

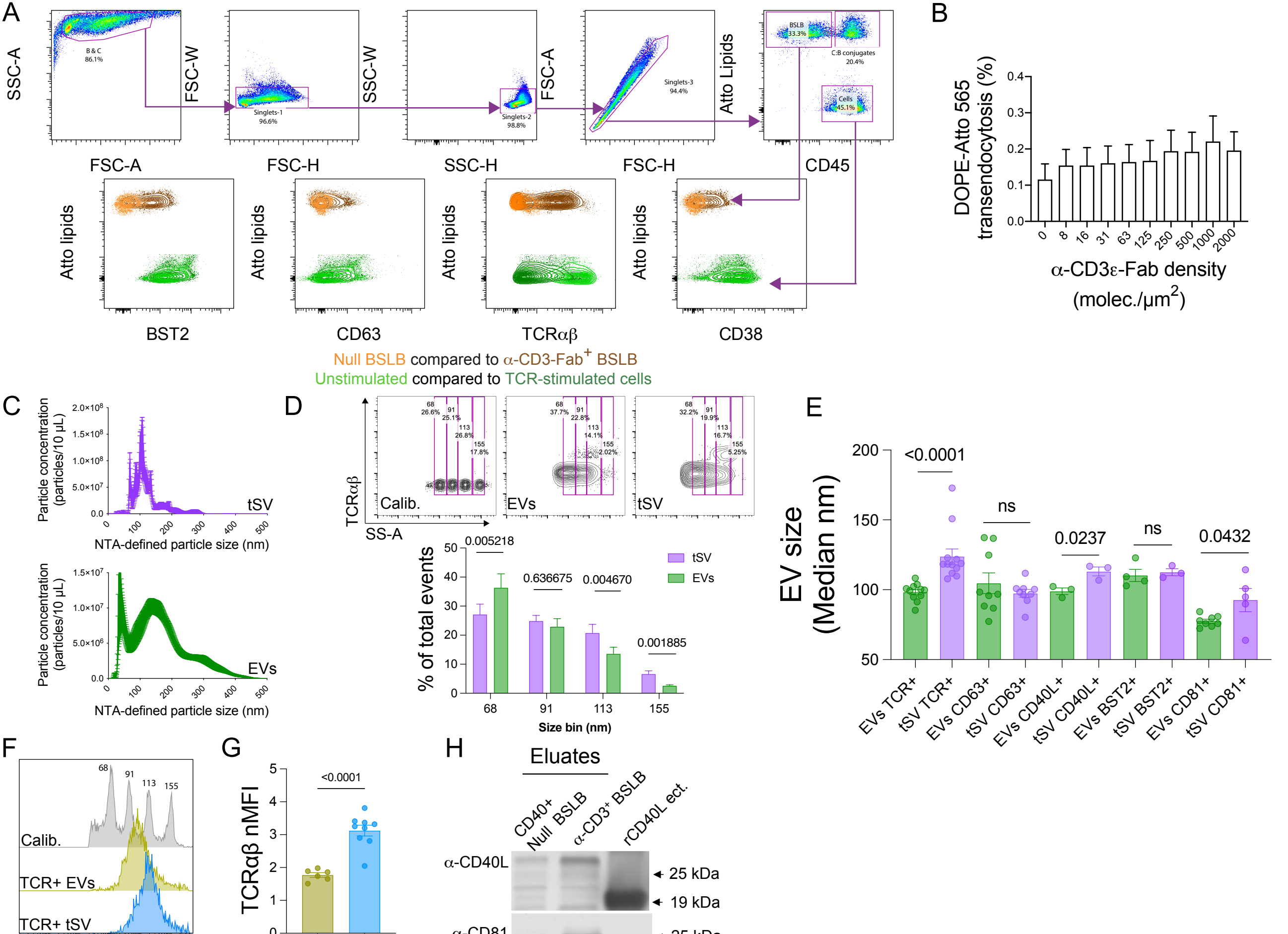


Supplemental Fig.1 related to Fig 1.

### Supplemental Figure 2

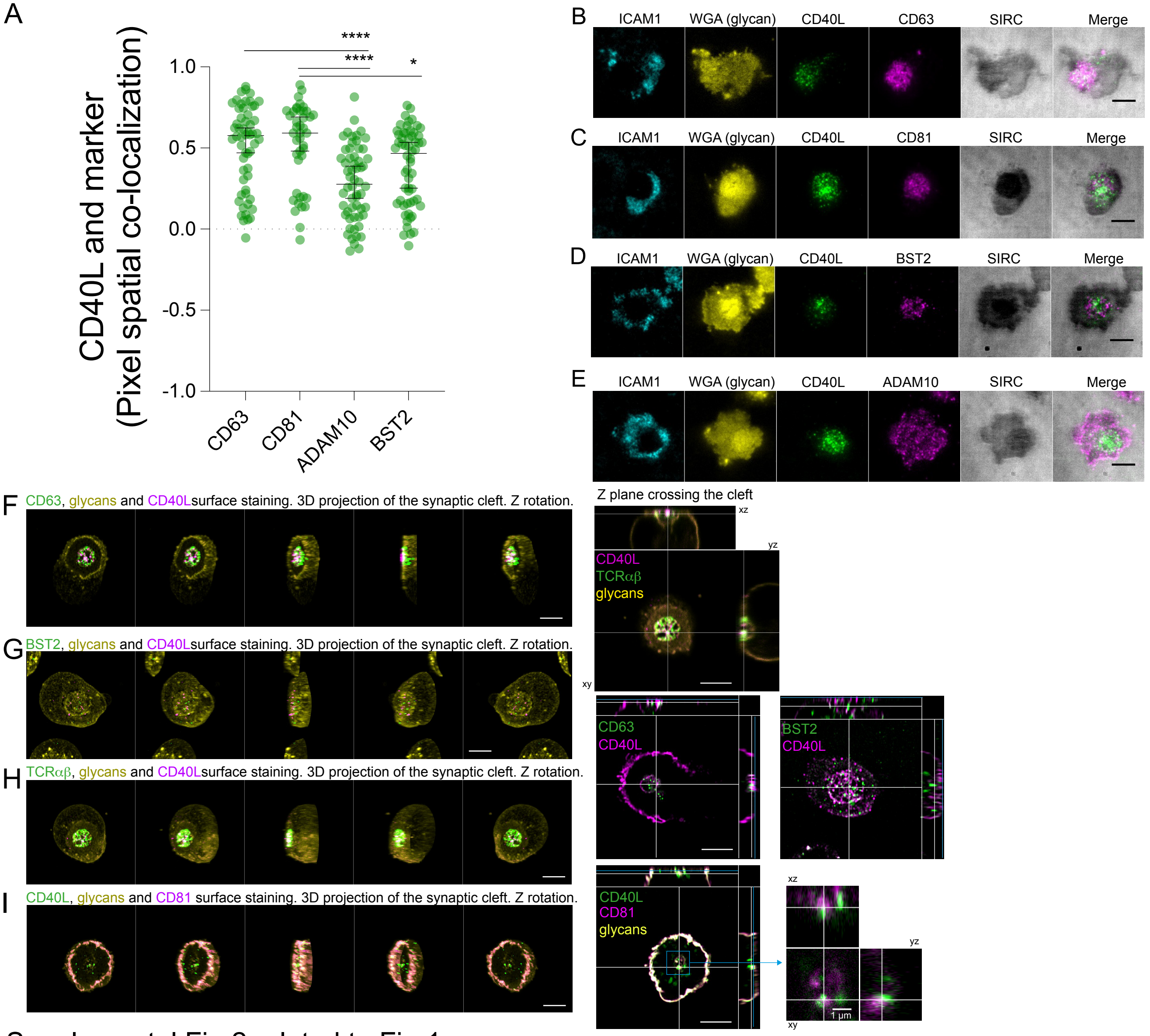


Supplemental Fig.2 related to Fig 1.

### Supplemental Figure 3

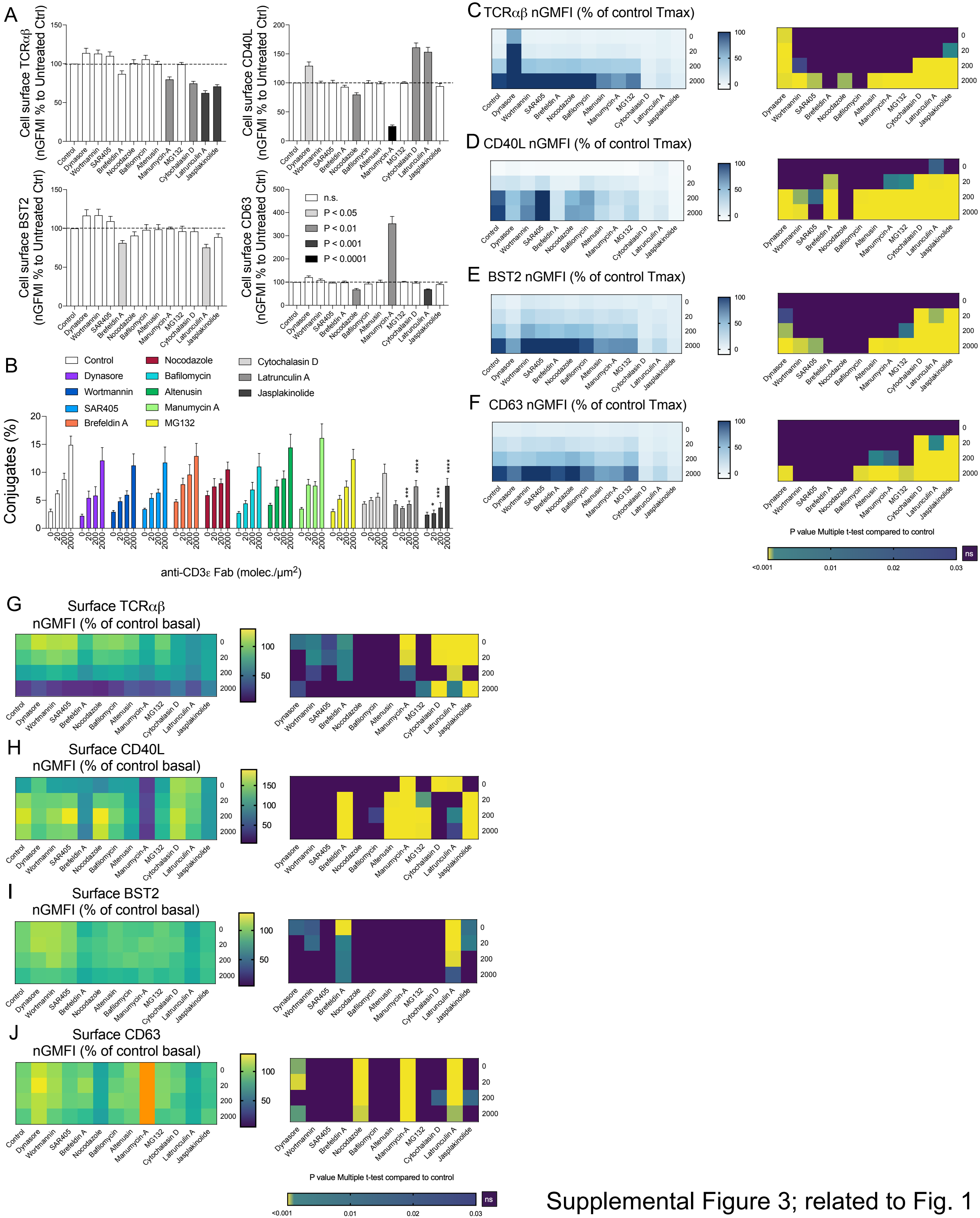


Supplemental Figure 3; related to Fig. 1

### Supplemental Figure 4

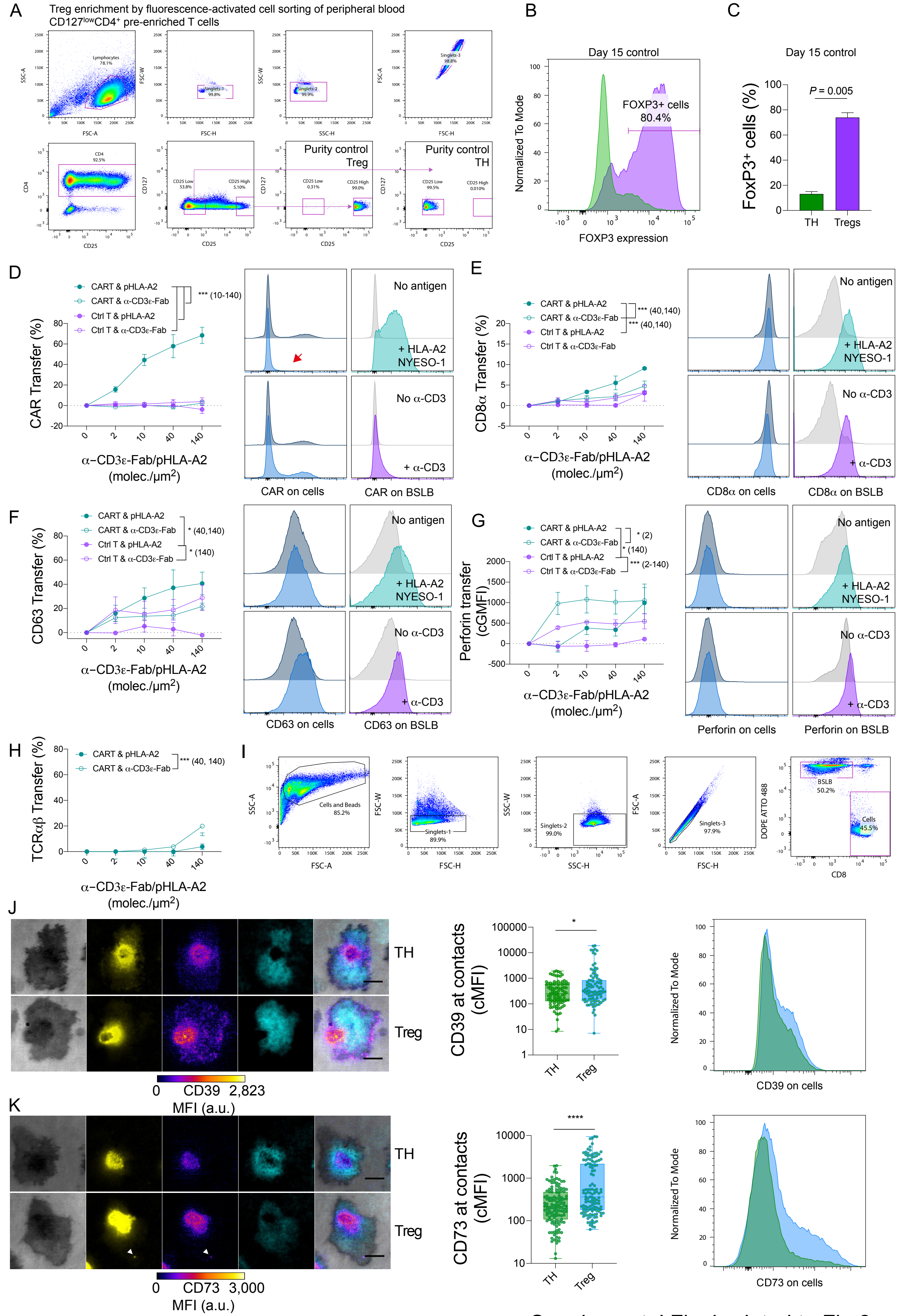


Supplemental Fig.4 related to Fig 2.

### Supplemental Figure 5

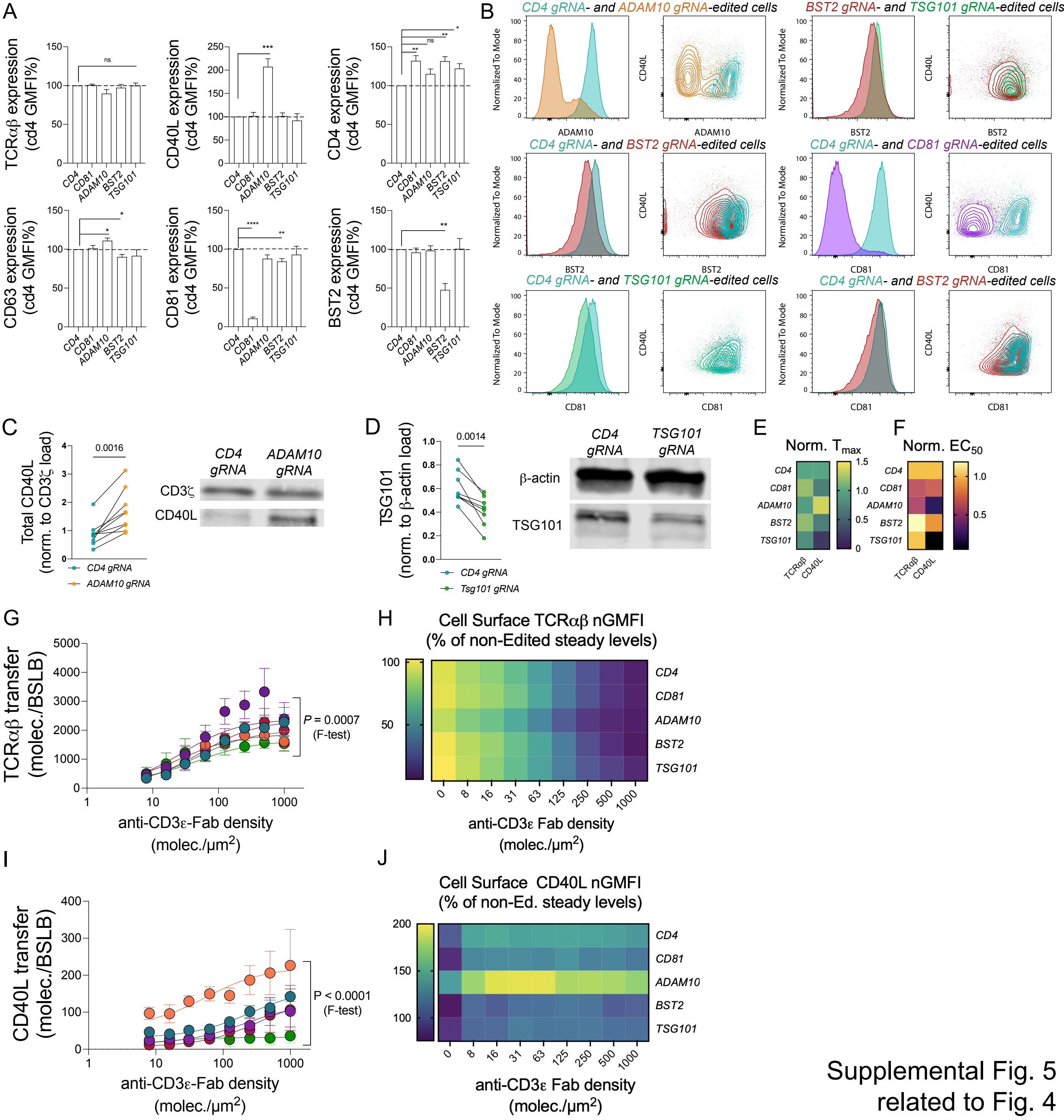


Supplemental Fig. 5 related to Fig. 4

### Supplemental Figure 6

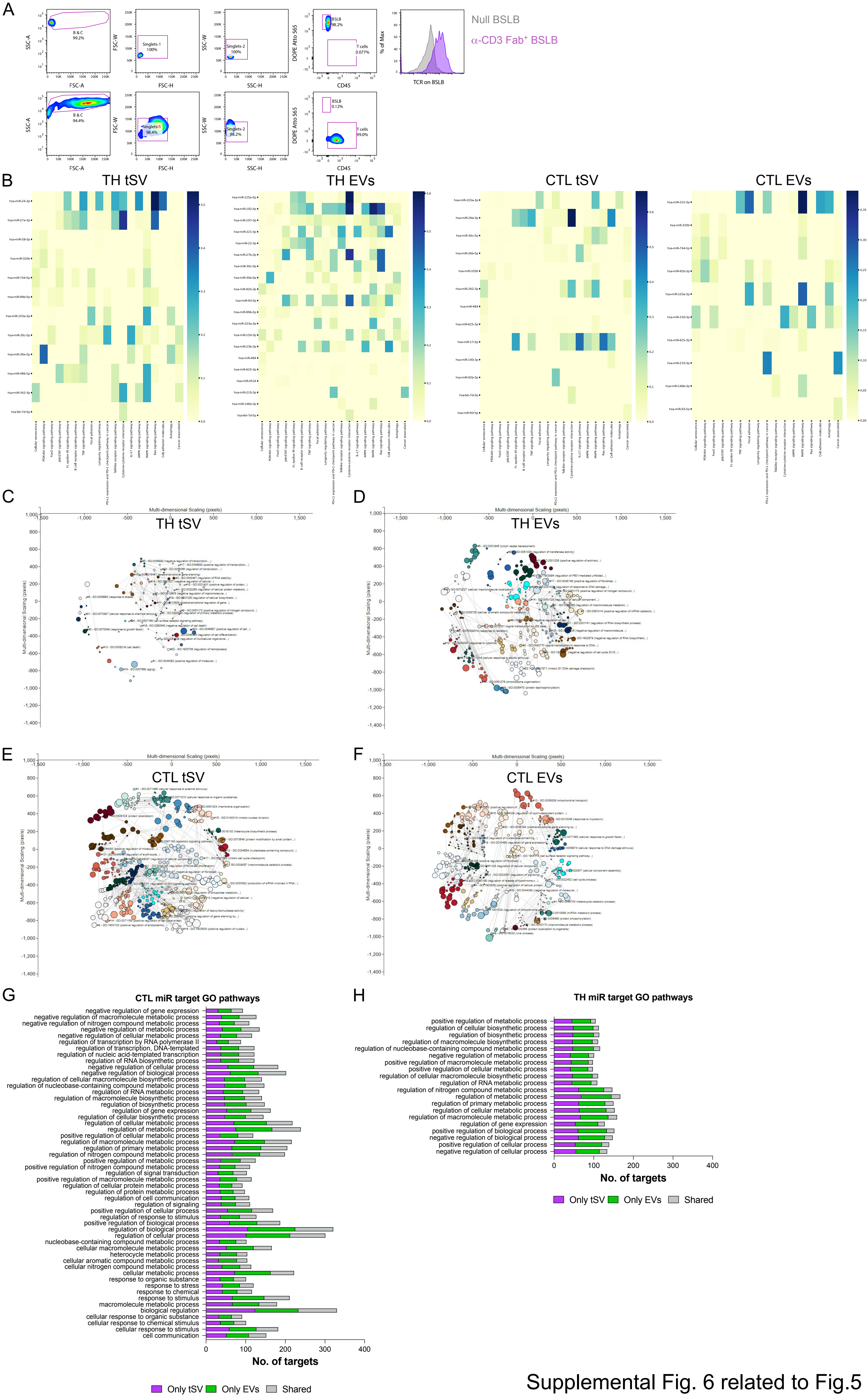


Supplemental Fig. 6 related to Fig.5

### Supplemental Figure 7

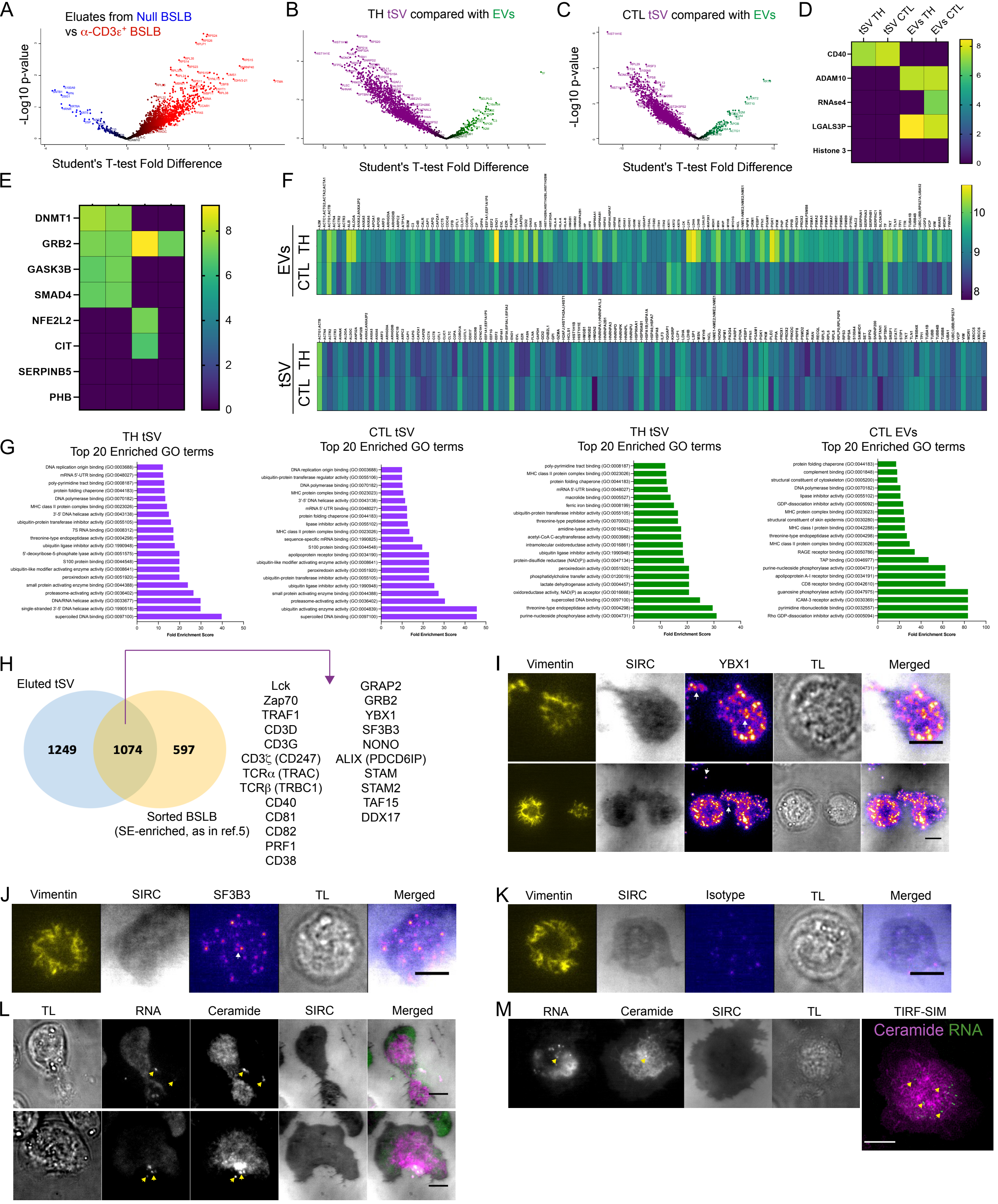


Supplemental Fig. 7 related to Fig. 5
